# Structure and function of one unclassified-class glutathione *S*-transferase in *Leptinotarsa decemlineata*

**DOI:** 10.1101/2021.09.29.462415

**Authors:** Yanjun Liu, Timothy Moural, Sonu Koirala BK, Jonathan Hernandez, Zhongjian Shen, Andrei Alyokhin, Fang Zhu

## Abstract

Arthropod Glutathione S-transferases (GSTs) constitute a large family of multifunctional enzymes that are mainly associated with xenobiotic or stress adaptation. GST-mediated xenobiotic adaptation is through direct metabolism or sequestration of xenobiotics, and/or indirectly by providing protection against oxidative stress induced by xenobiotic exposure. To date, the roles of GSTs in xenobiotic adaptation in the Colorado potato beetle (CPB), a notorious agriculture pest of plants within Solanaceae have not been well studied. Here, we functionally expressed and characterized an unclassified-class GST, LdGSTu1. The three-dimensional structure of the LdGSTu1 was solved with a resolution up to 1.8 Å by x-ray crystallography. Recombinant LdGSTu1 was used to determine enzyme activity and kinetic parameters using 1-chloro-2,4-dinitrobenzene (CDNB), GSH, p-nitrophenyl acetate (PNA) as substrates. The enzyme kinetic parameters and enzyme-substrate interaction studies demonstrated that LdGSTu1 could catalyze the conjugation of GSH to both CDNB and PNA, with a higher turnover number for CDNB than PNA. The LdGSTu1 enzyme inhibition assays demonstrated that the enzymatic conjugation of GSH to CDNB could be inhibited by multiple pesticides, suggesting a potential function of LdGSTu1 in xenobiotic adaptation.

## 1. Introduction

Glutathione *S*-transferases (GSTs) constitute a large superfamily of multifunctional enzymes that are ubiquitously present in prokaryotes and eukaryotes [1–4]. In general, GSTs catalyze the conjugation of the reduced glutathione (GSH) – a nucleophilic tripeptide comprised of three amino acids: cysteine, glutamic acid, and glycine – to a wide range of substrates that have an electrophilic carbon, nitrogen, or sulfur atom [5, 6]. The GST substrates can be natural or artificial compounds including cancer chemotherapeutic agents, carcinogens, pesticides, environmental pollutants, and byproducts of oxidative stress [4, 6]. In addition, GSTs are capable of binding numerous endogenous and exogenous compounds by non-catalytic interactions, which are associated with their functions in sequestration, storage, or transportation [3, 6, 7].

There are at least four major families of GSTs, namely cytosolic GSTs, mitochondrial GSTs, microsomal GSTs, and bacterial Fosfomycin-resistance proteins [3, 6, 8]. The first three families are present in both prokaryotes and eukaryotes. While the fourth family is only found in bacteria [8]. Two GST families, cytosolic GSTs and microsomal GSTs are identified in insects [1, 2]. The mitochondrial GSTs, also known as kappa class GSTs, are detected in mammalian mitochondria and peroxisomes but have not yet been identified in any insect species [1]. As soluble enzymes, insect cytosolic GSTs are divided into several classes based on their sequence similarities and structural properties: delta, epsilon, sigma, omega, zeta, theta, and unclassified classes [9–11]. Among these classes, delta and epsilon GSTs are insect-specific classes [12]. Insect cytosolic GSTs are biologically active as dimers with subunits ranging from 23-30 kDa in size. Each subunit consists of two domains joined by a variable linkage region [1, 3, 7, 13, 14]. The N-terminal domain constitutes a unique βαβαββα topology similar to the thioredoxin domain of many proteins that bind GSH or cysteine, suggesting an evolutionary relationship of cytosolic GSTs with glutaredoxins (GRXs) [1, 3, 14]. The N-terminal domain contains residues (e.g. cysteine, serine, or tyrosine) involved in binding and activating of GSH (the G-site) [1]. The C-terminal domain with a hydrophobic H-site shows a high level of diversity and is responsible for the interactions of GSTs with various electrophilic substrates [1, 7].

In insects, the functions of cytosolic GSTs are mainly associated with xenobiotic or stress adaptation [15]. For example, two delta class GSTs, BdGSTd1 and BdGSTd10, participate detoxification of malathion in the oriental fruit fly, *Bactrocera dorsalis* [16]. In *Drosophila melanogaster*, one delta GST (GSTD2) was proved to be involved in the detoxification of isothiocyanate [17]. In *Apis cerana cerana*, a sigma class GST (AccGSTS1) and a delta GST (AccGSTD) were suggested to have functions in cellular antioxidant defenses and honeybee survival [18, 19]. Another study found that a phytochemical induced epsilon GST (SlGSTe1) in the polyphagous insect pest *Sodoptera litura* could catalyze the conjugation of GSH with indole-3-carbinol, allyl-isothiocyanate, and xanthotoxin, suggesting its possible role in host plant adaptation [20]. In an Africa malaria vector, *A. funestus*, a single mutation in the binding pocket of GSTe2 coupled with increased transcription conferred high resistance to DDT and cross-resistance to pyrethroids [12].

Previous functional research on insect GSTs have mostly focused on delta, sigma, and epsilon classes. There are some unclassified GSTs sharing less than 40% amino acid identity with the other six insect GST classes, which have been temporarily designated unclassified (u) [1, 21]. The number of unclassified GSTs in each insect species is relatively small and the functions of these GSTs remain largely unclear [22]. For example, there are only three unclassified GSTs in the genome of *Anopheles gambiae* with 31 GST genes in total. Both *Tribolium castenaum* and *Bombyx mori* have two unclassified GST genes in their genomes. However, there is no unclassified GST identified in *D. melanogaster* and *A. mellifera* [22]. In the current study, we identified an unclassified GST gene, *LdGSTu1* from the Colorado potato beetle (CPB, *Leptinotarsa decemlineata* Say in Coleoptera which represents the most species-rich eukaryotic order, containing about half of the herbivorous insect species (>400,000) [23]. CPB is a global agriculture pest of the potato, *Solanum tuberosum* and other Solanaceae crops (e.g. tomato, eggplant, nightshade). This insect pest causes significant damage to potato crops by defoliation of plant leaves, which results in lose billions of dollars annually [24, 25]. One management strategy to control this problematic pest is the use of numerous pesticides. However, CPB is well recognized for its ability in rapidly adapting to various biotic and abiotic stresses, including almost all major classes of pesticides used for control [24, 26, 27]. The mechanisms of insects adaptation to pesticides and plant allelochemicals involves many aspects, including decreased penetration [28], target site insensitivity [29, 30], enhanced metabolic detoxification [31–33], increased excretion, sequestration as well as behavioral resistance [30, 34]. Among them, enhanced metabolic detoxification by cytochrome P450s, GSTs and other enzymes plays major roles in CPB xenobiotic adaptation [31, 35]. Uncovering function and structure of key enzymes in chemical adaptation pathways will help us understand mechanisms of stress adaptation in the global agricultural pest CPB [15, 31, 36, 37].

With about 30 crystal structures of insect GSTs having been solved, there is no crystal structure of a beetle GST available (Table S1). Here, we solved the LdGSTu1 co-crystal structure with its nucleophilic substrate GSH by using X-ray diffraction. The three-dimensional structure of the LdGSTu1 was solved with a resolution up to 1.8 Å by x-ray crystallography. A typical GST global fold and an active site composed of two substrate binding sites, the “G-site” and the “H-site” were identified. The LdGSTu1 enzyme kinetic parameters and enzyme-substrate interaction studies demonstrated that the conjugation of GSH to CDNB could be inhibited by multiple pesticides, suggesting a potential function of LdGSTu1 in pesticide adaptation.

## 2. Materials and methods

### 2.1. Insects

The susceptible CPB was purchased from French Ag Research, Inc. (Lamberton, MN), originally collected from Long Island in 2003 and reared under laboratory conditions without exposure to any pesticides. The insecticide resistant CPB population was collected from commercial potato fields in Presque Isle, Maine. Both populations were reared on Red Norland potato plants in several BugDorm insect cages (MegaView Science Education Services Co., Ltd.) at 25 ± 5 °C under a light:dark regimen of 16:8 h in a Penn State facility greenhouse. New plants were provided once every week. Eggs were collected each day and stored in petri dishes kept at 25 ± 1°C, RH of 70%, and L:D=16:8. After emergence larva were fed on fresh potato leaves until reaching the 2^nd^ instar when they were transferred back to greenhouse rearing cages.

### 2.2. LdGSTu1 cloning, bioinformatics, and phylogenetic analysis

The LdGSTu1 cloning was performed with a ligation-independent cloning strategy following a previous protocol [38]. Briefly, the full-length LdGSTu1 was amplified from resistant CPB cDNA using PCR with the primers containing ligation independent cloning F and R sites (Table S2), T4 polymerase treated, and then annealed with T4 polymerase treated pET-9Bc vector. The pET C-terminal TEV His6 cloning vector with BioBrick polycistronic restriction sites (9Bc) was a gift from Scott Gradia (Addgene plasmid #48285; http://n2t.net/addgene:48285; RRID:Addgene 48285). Then the products were transformed into DH 5α competent cells. Positive colonies were verified using T7 primers, then cultured in liquid LB overnight at 37 ℃. The plasmids were extracted and identified, and sequenced by Functional Biosciences, Inc. The cloned sequences were submitted to the website of National Center for Biotechnology Information (NCBI) (https://www.ncbi.nlm.nih.gov/). The conserved domains were detected using bioinformatics tools on the NCBI server. The theoretical isoelectric point (pI) and molecular weight (MW) were computed using the Compute pI/Mw tool (https://web.expasy.org/compute_pi/). To classify the GST gene, the phylogenetic tree was constructed with Muscle and MEGA X using the maximum likelihood, LG model, gamma distributed method with 1000 bootstrap replicates [39, 40]. The available amino acid sequences of GSTs used in the phylogenetic analysis were downloaded from the NCBI database [41]. Multiple alignment analysis was also conducted with several GTSs from different insects by DANMAN v. 6.03 (Lynnon BioSoft, Vaudreuil, Quebec, Canada).

### 2.3. LdGSTu1 protein expression and purification

The pET-9Bc-LdGSTu1 plasmids were transformed into Rosetta^TM^ II (DE3) pLysS, positive colonies were verified with PCR. The successful inserts were grown in 50 mL LB cultures at 37 °C in incubator (Thermo Scientific MaxQ 6000 Incubated Stackable Floor Shaker) for induction testing. Cell stocks positive for LdGSTu1 expression were frozen at −80°C for later using. For expression, overnight 50 mL cultures grown at 37 °C in terrific broth with 1x ampicillin and chloramphenicol, and then used to inoculate a 1.2 L Terrific Broth (TB) culture, incubated at 37 °C until OD_600_ reached 0.4 ∼ 0.6. Then the culture was cooled and induced with 0.5 mM IPTG and incubated at 20 °C for an additional 20 hours. Cells were harvested by centrifugation at 4,000 rpm and 4 °C to acquire cell pellet in a tabletop centrifuge (Thermo Sorvall Legend XTR refrigerated centrifuge). The cells were lysed with buffer containing 50 mM NaPi, 300mM NaCl, 20 mM Imidazole, and 1 mM PMSF (pH: 7.6) for 5 cycles of 30 seconds at 70% power 5-7 times using a sonicator (Branson Digital Sonifier SFX 150) on ice. After cell lysis, the homogenate was centrifuged at 18,000g to separate soluble protein from the insoluble fractions. The soluble fraction was added to a Ni-NTA column. The Ni-NTA column was washed with 50 mM NaPi, 300mM NaCl, 20 mM Imidazole (pH: 7.6). Next, LdGSTu1 protein was eluted with buffer containing 20 mM NaPi, 300 mM NaCl, and 250 mM Imidazole (pH: 8.0). Protein was then 100 x fold buffer exchanged into buffer containing 5 mM NaPi, 5 mM HEPES (pH: 7.2). Next, protein was further purified by a Hydroxyapatite column (HA) connected to an NGC Medium-Pressure Liquid Chromatography System (BIO-RAD). A gradient form 5 mM NaPi, 5 mM HEPES (pH: 7.2) to 500 mM NaPi (pH: 7.2) was used to wash and elute LdGSTu1. Fractions containing LdGSTu1 were visualized with SDS-PAGE, combined, and concentrated. Then concentrated fractions were applied to SEC Enrich 650 (BioRad) connected to the NGC. Size exclusion buffer was 20 mM HEPES, 1 mM DTT, and 1 mM EDTA, 20 mM GSH pH 7.2. Fractions were checked with SDS-PAGE. Purified LdGSTu1 protein concentrations were calculated using the Bradford assay or UV280 methods on NanoDrop One (Thermo Scientific) and Spark multi-mode plate reader (Tecan). Protein was concentrated and used directly for crystallization or buffer exchanged for enzyme assays.

### 2.4. Crystal data collection, refinement, and structural analysis

LdGSTu1 crystals were grown by sitting drop vapor diffusion at 18 °C. High purity LdGSTu1 at 20mg/mL was mixed 1:1 with reservoir solution in sitting drop well and incubated against mother liquor reservoir solution (100 mM MES pH 6.5, 100mM NaCl, 25% PEG 4K). LdGSTu1 crystal data were collected at the Macromolecular X-ray science at the Cornell High Energy Synchrotron Source (MacCHESS) beamline 7B2. The software package XDS was used for data processing [42]. Phasing was done by using BmGSTu1 (PDB: 5ZFG) as a search model in PHENIX Phaser [43]. Refinement and model building were performed by using PHENIX and Coot [43–45]. Search models for molecular replacement were identified by a NCBI blastp with the LdGSTu1 amino acid sequence search against the Protein Data Bank (PDB) database [46, 47]. Structural analysis and figures of LdGSTu1 were conducted by using UCSF Chimera, UCSF Chimera X, and Coot [48–50]. The coordinates and structure factors for the final model of LdGSTu1 and GSH was deposited in the PDB under accession code 7RKA.

### 2.5. Enzyme assay

The kinetic analysis of LdGSTu1 was conducted by steady state with varied concentrations of substrates 1-chloro-2,4-dinitrobenzene (CDNB) from 0.05 mM to 3 mM and p-nitrophenyl acetate (PNA) from 0.2 mM to 3.2 mM, while holding the GSH concentration constant at 5 mM, and for varied concentrations of GSH at 0.125 mM to 5mM while holding CDNB at a constant concentration of 2 mM. The reaction buffer was 100 mM KPi (pH: 6.5). Reactions were carried out in 96-well UV-Star^®^ microplates (Greiner Bio-One). Assays were run on a Tecan Spark^®^ multi-mode plate reader in the kinetic mode, for a continuous read assay for 3 mins. Product concentrations were calculated by path-length corrected molar attenuation coefficient from Habig et al. [51, 52]. Kinetic parameters and plots were calculated and generated with GraphPad Prism.

### 2.6. Enzyme inhibition assay

Residual enzyme activity in the presence of inhibitors was measured by individually incubating LdGSTu1 (50 μg) with inhibitors ethacrynic acid (EA) and carbaryl, diazinon, imidacloprid, acetamiprid, chlorpyrifos, and thiamethoxam at 40 µM, 200 µM, 1mM, and 5 mM for 10 minutes at 30°C followed by addition of GSH (0.5 mM) and CDNB (0.5 mM), for total reaction volume of 200 μL in 96-well Greiner Bio-One UV-STAR^®^ microplates. After addition of GSH and CDNB the change in absorbance at a wavelength of 340 nm was immediately measured on a TECAN Spark^®^ multimode plate reader at 30°C in kinetic mode for 70 secs with 10 second reads. at. All inhibition and control reactions were run in triplicate. Residual activity was calculated as precent of enzyme activity retained in reaction in presence of inhibitor relative to control reaction without inhibitor and an equivalent amount of acetone. Additionally, controls without enzyme were run to control for non-enzymatic reaction contribution. Inhibitor stock solutions were prepared in acetone. For carbaryl reactions were only run at 40 µM, 200 µM, and 1mM due to insolubility for the reaction conditions at 5 mM. Also, EA reactions were only run at 40 µM, 200 µM, and 1mM due to having reached 100% inhibition at 1mM. The reaction buffer was 100 mM KPi at a pH of 6.5.

### 2.7. Docking of LdGSTu1 crystal structure with xenobiotics

The LdGSTu1 crystal structure was prepared for docking studies with Molecular Operating Environment (MOE), 2020.9 [53]. The refined LdGSTu1 crystal structure was prepared with the Structure Preparation, Protonate 3D, and Partial Charge applications. The molecular mechanics forcefield used was Amber10:EHT. The ligand structures of CDNB, EA and carbaryl, diazinon, imidacloprid, acetamiprid, chlorpyrifos, and thiamethoxam were downloaded from PubChem^®^ database in SDF format [54]. Ligands were docked with the MOE Dock application using the prepared LdGSTu1 chain A structure containing complexed GSH as the receptor and the substrate binding pocket selected as ligand binding site. The method parameters selected were Triangle Matcher for placement with scoring function London dG, 30 poses, and refinement of Induced Fit with scoring function Affinity dG, 5 poses. Ligand docking runs gave 5 final poses for each ligand with favorable binding energy. The highest ranked docked ligand poses were selected for further analysis. Figure with docked ligands was generated with ChimerX [55].

### 2.8. RNA extraction, cDNA synthesis and qRT-PCR analysis

Total RNA was isolated from insect samples using TRIzol reagent (Invitrogen, Maryland, USA). The total RNA was treated with DNase I (Ambion Inc., Austin, TX, United States) to remove contaminating genomic DNA. Approximately 3 *μ*g of DNase I-treated RNA was reverse transcribed to cDNA using 10 *μ*M of oligo (dT) primer and M-MLV reverse transcriptase (Promega, Madison, WI USA) in a 20 *μ*L reaction. NanoDrop 2000 spectrophotometer (Thermo Fisher Scientific, USA) was used to perform spectroscopic quantification. The *ef1α* and *rpl4* were used as reference genes for the qRT-PCR [31]. PCR conditions included 3 min at 95°C followed by 39 cycles of 10 s at 95°C, and 55°C for 30 s. qRT-PCR was conducted with 1 *μ*L cDNA, 5 *μ*L FastStart SYBR Green Master (Roche Diagnostics, Indianapolis, IN USAl), 0.4 *μ*L qRT-PCR primers (Table S2), and 3.6 *μ*L ddH_2_O in a 10 *μ*L total reaction volume using Bio-Rad CFX Connect™ Real-Time PCR System (Bio-Rad, CA, USA). The 2^-ΔΔCt^ method was used for the quantitative analysis. Three biological replications were conducted independently.

### 2.9. Developmental and spatial expression of LdGSTu1

The different developmental stages including eggs (first day and fourth day); larvae (1^st^-4^th^ instar); pupa; female and male adults were collected. At least 10 individuals were used for each stage. Different tissues including head (with antenna), midgut, malpighian tubule, fat body, and ovary from adult were dissected in ice-cold 1x phosphate-buffered saline (PBS) solution. Each tissue collected from at least 20 adults (sex ratio = 1:1) was pooled as one sample. Three independent replicates were performed.

### 2.10. Statistical analyses

All statistical analyses were conducted using SPSS 20.0 (SPSS Inc., Chicago, USA). Data were expressed as the mean ± standard error (SE) of three independent replicates. Differences in gene expression between two experimental treatments were analyzed using independent Student’s *t*-test. Differences among multiple treatments were analyzed by one-way ANOVA, followed by a Tukey’s HSD multiple comparison test (*P* < 0.05). In Student’s *t*-test, significance levels were denoted by * (0.01 < *P* < 0.05) and ** (*P* < 0.01).

## 3. Results

### 3.1. Phylogenetic relationship of LdGSTu1 with other insect GSTs

The *LdGSTu1* gene was cloned from the *L. decemlineata* susceptible and resistant strains and shared 100% sequence similarity with the gene XP_023027125.1 in NCBI database. Sequence analysis showed that the ORF was 693 bp, encoding a deduced polypeptide of 230 amino acids. The predicted molecular weight of LdGSTu1 was 26.4 kDa and isoelectric point was 5.36. The phylogenetic tree was constructed by the maximum likelihood method using the deduced amino acid sequences to investigate the evolutionary relationships of LdGSTu1 and 31 GSTs from *L. decemlineata* and other insect species (Fig. 1, Table S3). Phylogenetic tree showed that the GSTs from the same class were grouped together (Fig. 1). As expected, LdGSTu1 was clustered in the unclassified clade with other five unclassified GSTs identified from *Anoplophora glabripennis*, *Drosophila mauritiana*, *D. mojavensis*, *Bombyx mori* and *Sitophilus oryzae* (Fig. 1, Table S3). LdGSTu1 originated from the same evolutionary root with SoGST1-X2 from *Sitophilus oryzae* with the bootstrap value of 75 (Fig. 1).

**Fig. 1.**
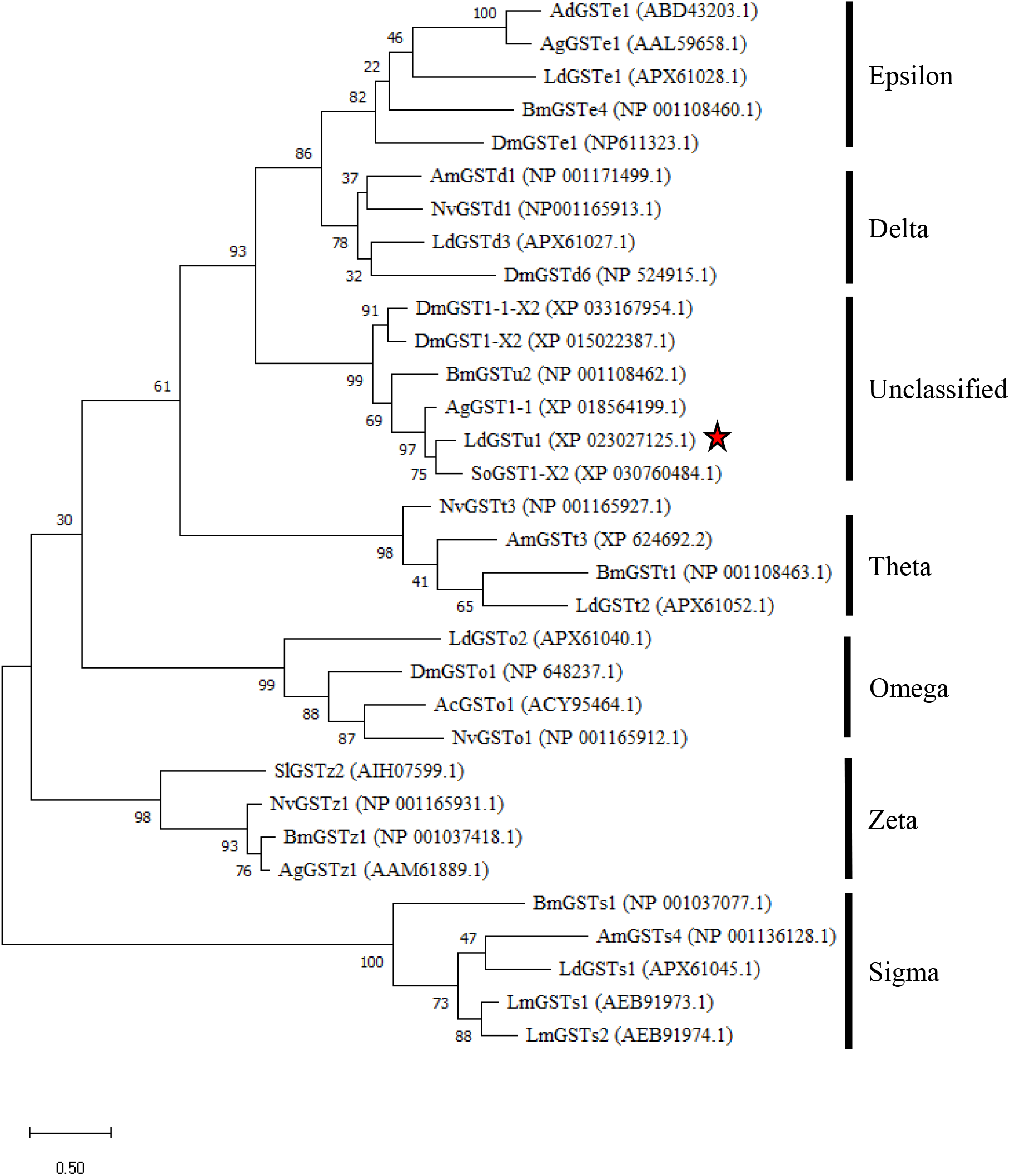
Phylogenetic analysis. Phylogenetic analysis of LdGSTu1 with homologs in other insects. Ac, Anopheles cracens; Ad, Anopheles dirus; Ag, Anopheles gambiae/Anoplophora glabripennis; Am, Apis mellifera; Bm, Bombyx mori; Dm, Drosophila mauritiana/Drosophila mojavensis/Drosophila melanogaster; Ld, Leptinotarsa decemlineata; Lm, Locusta migratoria; Nv, Nasonia vitripennis; Sl, Spodoptera litura; So, Sitophilus oryzae. The red star indicates LdGSTu1.

### 3.2. X-ray crystal structure of LdGSTu1 in complex with GSH

LdGSTu1 crystalized in space group P2 with a unit cell of a = 58.45, b = 46.44, and c = 87.19. The LdGSTu1 structure was refined to a resolution of 1.80 Å. Two monomers of LdGSTu1 were in the crystal asymmetric unit. One monomer, Chain A exhibited glutathione (GSH) bound to the active site (Fig. 2). Whereas the other monomer (Chain B) had no bound GSH molecule (Fig. S1). Data collection and refinement statistics are listed in Table 1.

**Fig. 2.**
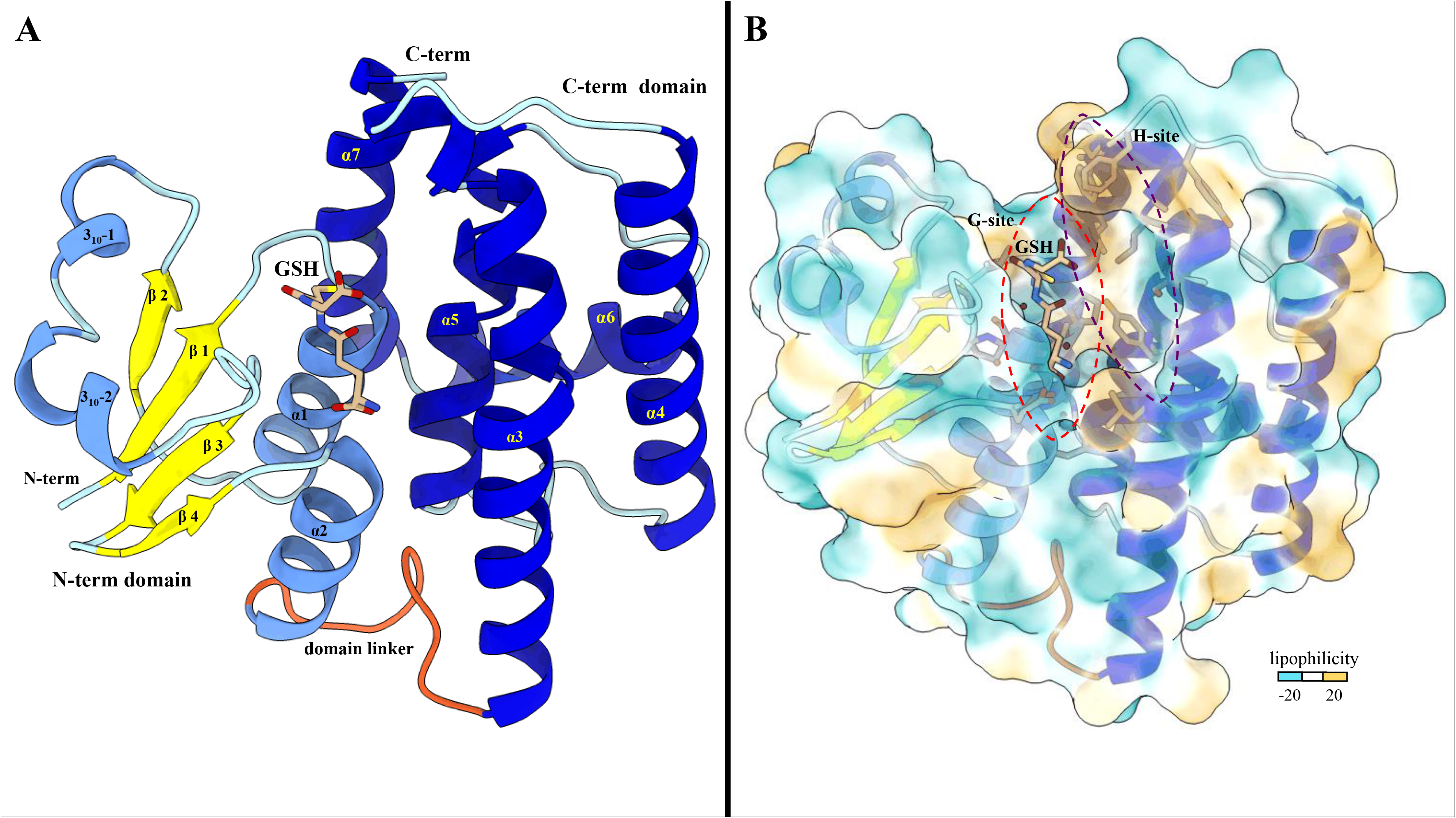
Global structure of LdGSTu1. (A) Ribbon diagram of LdGSTu1 with GSH bound in the active site, showing secondary structural elements and the N-term and C-term domain structures. The N-term domain exhibits the thioredoxin-like fold (residues 1-77) and was complexed with GSH in chain A of the crystal structure. Helices are depicted in cornflower blue, coils in powder blue, and β-strands in yellow. The linker coil (residues 78-88) connecting the polypeptide of the N-term domain to the polypeptide of the C-term domain is depicted in orange. The C-term domain is mainly helical in nature, consisting of five α-helices (medium blue) with coils depicted in powder blue. (B) MLP surface diagram of LdGSTu1 monomer with bound GSH in active site. The color scale is cyan for most hydrophilic and passes through white to golden rod for most hydrophobic. In the active site the most hydrophilic location was the G-site with GSH bound and the most hydrophobic region of the active site was the H-site on the C-term side of the active site.

**Table 1.** Data collection and refinement.

| **Parameter** | **LdGSTu1 complex with GSH** |
| --- | --- |
| Wavelength (Å) | 0.9686 |
| Resolution range (Å) | 29.22 - 1.8 (1.864 - 1.8) |
| Space group | P 1 2 1 |
| Unit cell | a = 58.452 b = 46.44 c = 87.19<br>$\alpha = 90 \beta = 91.35 \gamma = 90$ |
| Total reflections | 135715 (7106) |
| Unique reflections | 42482 (3476) |
| Multiplicity | 3.2 (2.0) |
| Completeness (%) | 97.12 (79.63) |
| Mean I/sigma(I) | 16.28 (2.71) |
| R-merge | 0.05293 (0.3295) |
| R-meas | 0.05456 (0.5671) |
| R-pim | 0.0286 (0.2128) |
| CC1/2 | 0.999 (0.888) |
| CC* | 1 (0.97) |
| **Reflections used in refinement** | 42476 (3476) |
| Reflections used for R-free | 1741 (133) |
| R-work | 0.1599 (0.2253) |
| R-free | 0.1835 (0.2637) |
| CC (work) | 0.965 (0.915) |
| CC (free) | 0.955 (0.876) |
| **Number of non-hydrogen atoms** | 3753 |
| Macromolecules | 3416 |
| Ligands | 20 |
| Solvent | 317 |
| Protein residues | 422 |
| **Average B-factor** | 27.49 |
| Macromolecules | 26.7 |
| Ligands | 41.50 |
| Solvent | 35.2 |

#### Overall Structure of LdGSTu1

A NCBI blastp search with the LdGSTu1 sequence revealed that the highest identity matches with the PDB published unclassified GST, BmGSTu2 (PDB: 5ZFG) at a sequence identity of 60.43% [46, 47, 56]. In regards to insect GST classified classes (Delta, Epsilon, Omega, Sigma, Theta, and Zeta), LdGSTu1 exhibits the highest precent identities to Delta class, 40.38%, 39.62%, and 38.21% with AgGSTD 1-6 (PDB: 1PN9), AcGSTD 1-3 (PDB: 1JLV), and NlGSTD (PDB: 3WYW) [57–59], respectively.

The global fold of LdGSTu1 is representative of the “GST fold” similar to previously published structures of insect GSTs (Fig. 2A) [56–59]. LdGSTu1 consists of two domains, the N-term domain and the C-term domain connected by a linker region coil. The N-term domain comprises four β-strands, two α-helices, and two 3_10_-helices. The secondary structural elements of the N-term domain are ordered starting at the N-terminus with β1(residues 3-7), followed by α1 (residues12-23), β2 (residues 29-32), 3_10_-1 (residues 36-40), 3_10_-2 (residues 43-48), β3 (residues 56-59), β4 (residues 62-64), and α2 (residues 67-77). Then the linker coil (residues 78-88) connects the N-term domain to the C-term domain. The C-term domain consists of five helices, α3 (residues 89-119), α4 (residues 126-146), α5 (residues 158-173), α6 (residues 181-193), and α7 (residues 195-211) (Fig. 2A and Fig. 3). A substrate binding pocket is located between the N-term domain and the C-term domain. GSTs have been previously described as having two binding sites within the substrate binding pocket, the “G-site” which binds GSH and “H-site” which binds a hydrophobic co-substrate [58]. A molecular lipophilicity potential surface reveals that the LdGSTu1 substrate binding pocket has a more hydrophilic region on the N-term domain side of the binding pocket, which is bound with GSH and makes up the “G-site”, and a more lipophilic region in the substrate binding pocket adjacent to the bound GSH on the C-term domain side of the pocket making up the “H-site” (Fig. 2B).

**Fig. 3.**
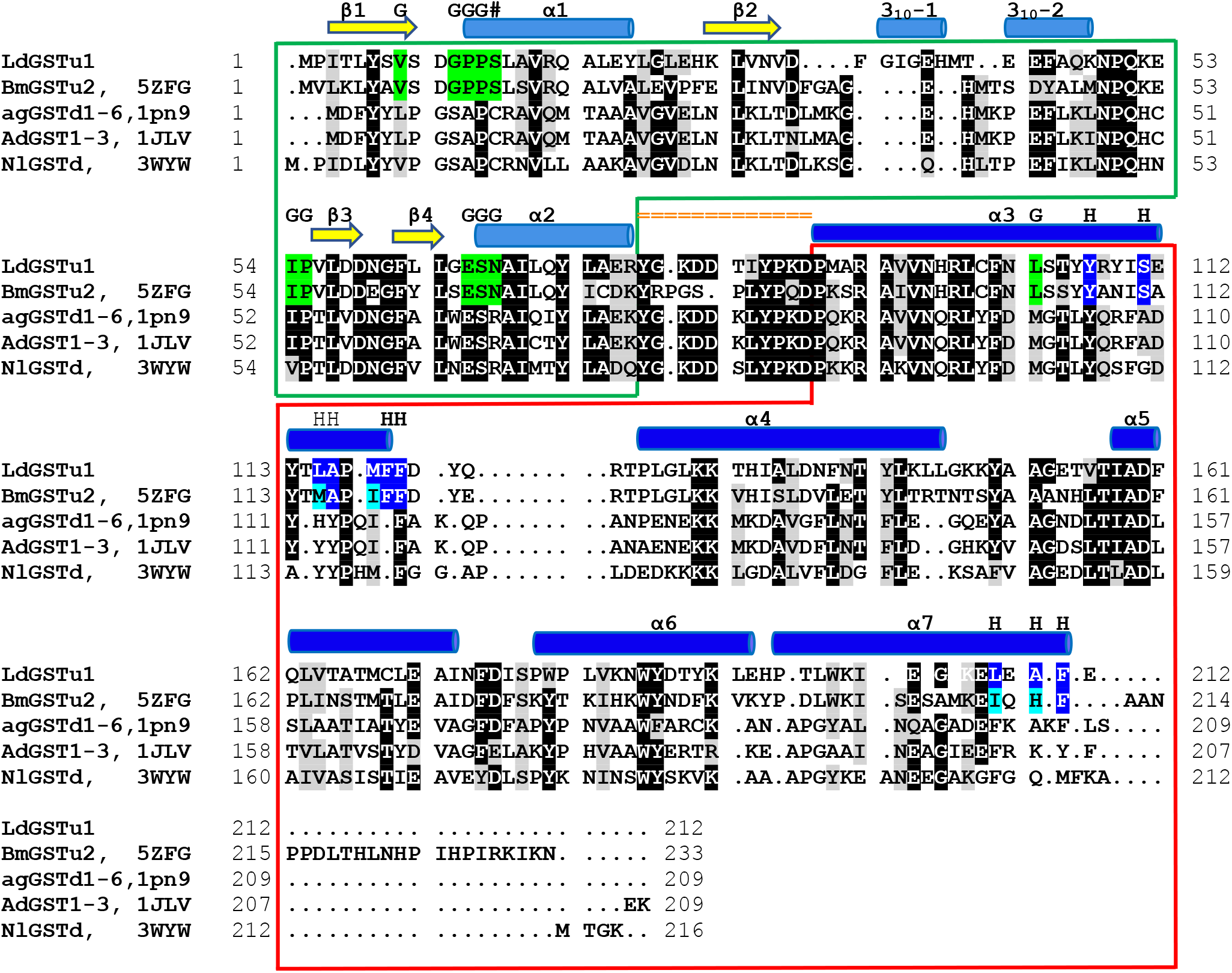
Structural sequence alignment of LdGSTu1 with highest sequence identity insect GSTs deposited in PDB. Chimera was used to superpose LdGSTu1 crystal structure with 5ZFG, 1PN9, 1JLV, and 3WYW to generate a multiple structure sequence alignment. Sequence amino acids are highlighted black for 80% identity and gray for 80% similarity. Secondary structural element positions are mapped and shown with yellow arrows for β-strands, and blue cylinders for helices. Domain distribution is depicted as boxed green for the N-term domain and boxed in red for the C-term domain. The linker region is mapped with orange equal signs. G-site residues are highlighted green and marked with a G. H-site residues are highlighted blue and marked with an H.

#### Active site of LdGSTu1

The active site of LdGSTu1 is formed by a pocket consisting of a “G-site” which binds GSH and an “H-site” which binds a hydrophobic substrate, consistent with previously insect GST studies (Fig. 2B) [56, 58]. The “G-site” is depicted in Fig. 4A with all side chains of all amino acids within 4.5 Å. The GSH is bound to LdGSTu1 via an extensive hydrogen bonding network that includes amino acid side chains, backbone carbonyls, backbone amides and crystallographic water. Ile54 and Ser64 form direct hydrogen bonding interactions with GSH (Fig. 4A). Ser14, Pro55, Asn68, and Glu66 form hydrogen bonds bridged by crystallographic water molecules with GSH. The presumptive catalytic Ser14 formed a hydrogen bond with the thiol group of GSH bridged via a water molecule. Additionally, the side chain OE1 and OE2 atoms of Glu66 formed an ionic interaction with N1 atom of the bound GSH. The “H-site” of LdGSTu1 is largely hydrophobic (Fig. 2B) and consists of Tyr107, Ser111, Leu115, Ala116, Phe119, Phe120, Leu208, Ala210, and Phe211 (Fig. 4B).

**Fig. 4.**
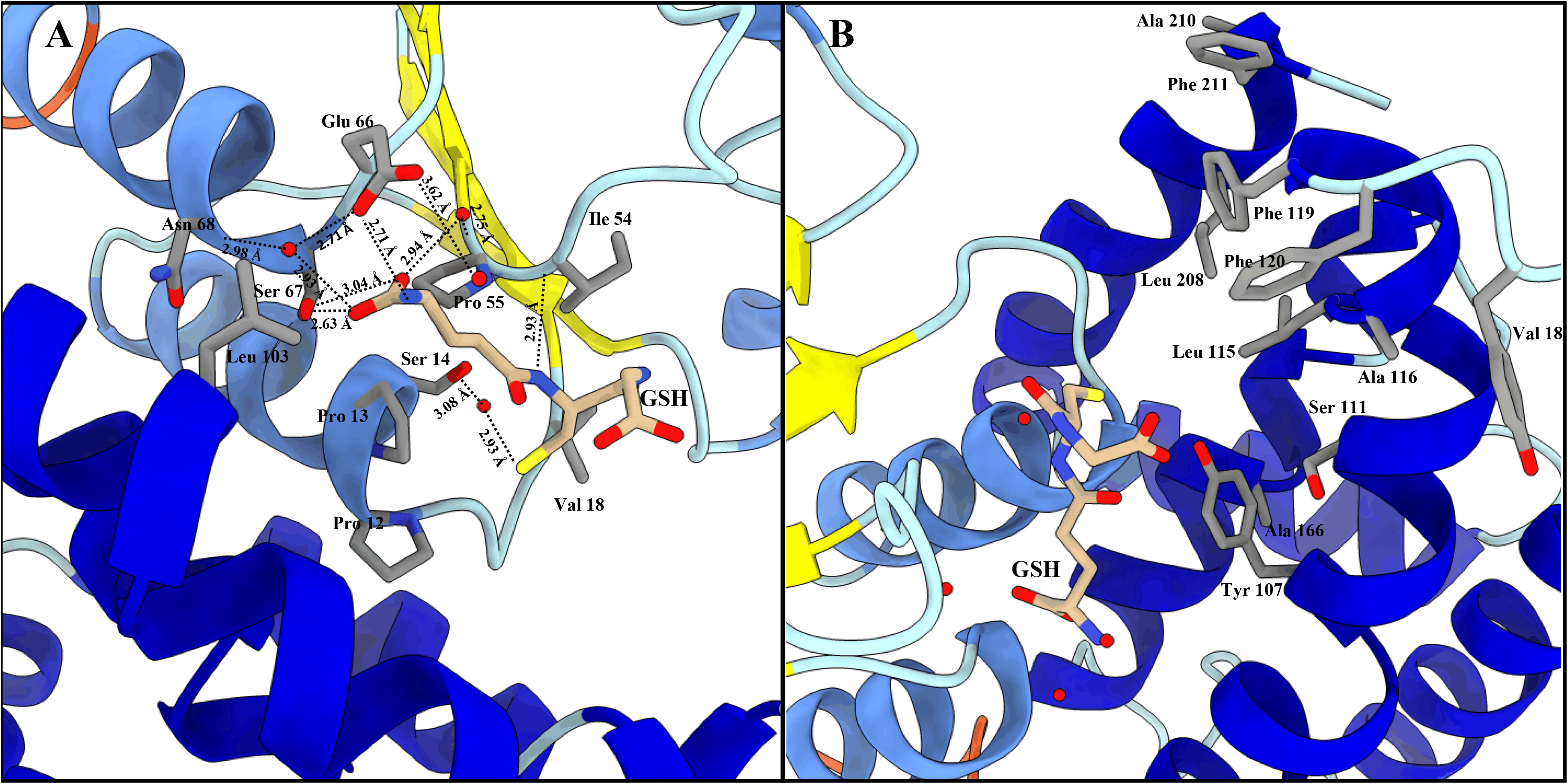
LdGSTu1 active site with GSH. (A) Zoomed in view with substrate GSH bound to LdGSTu1 active site. An extensive hydrogen bonding network was established between LdGSTu1, crystallographic water molecules and the GSH substrate. Hydrogen bond lengths between LdGSTu1 residues, waters, and GSH are shown with dashed lines and given bond lengths are given in angstrom. (B) Adjacent to the bound GSH, is the H-site located on the C-term domain side of the active site, side chains of the amino acids making the presumptive hydrophobic substrate binding site are shown in elemental color scheme.

### 3.3. Enzymatic properties of LdGSTu1

The kinetic analysis of LdGSTu1 was conducted by steady state with varied concentrations of substrates CDNB and PNA while holding the GSH concentration constant, and for varied concentrations of GSH while holding CDNB at a constant concentration. Michaelis-Menten plots were generated, and curve fit by nonlinear regression with GraphPad Prism (Fig. 5). Kinetic parameter values were found to be: Vmax values were 78.2 ± 3.46 μM/min, 60.9 ± 3.49 μM/min, 13.5 ± 2.13 μM/min; the Km values were 0.689 ± 0.118 mM, 0.542 ± 0.088 mM, 1.830 ± 0.572 mM; the k_cat_ were 44.0 ± 1.95 min^-1^, 34.1 ± 0.63 min^-1^, 7.7 ± 1.21 min^-1^; and k_cat_/K_m_ values were 63.8 mM/min, 62.9 mM/min, 4.2 mM/min; for GSH, CDNB, and PNA, respectively (Table 2). However, LdGSTu1 was not active against 4-hydroxynonenal (HNE) and trans-2-hexenal (T2H) (Table 2).

**Fig. 5.**
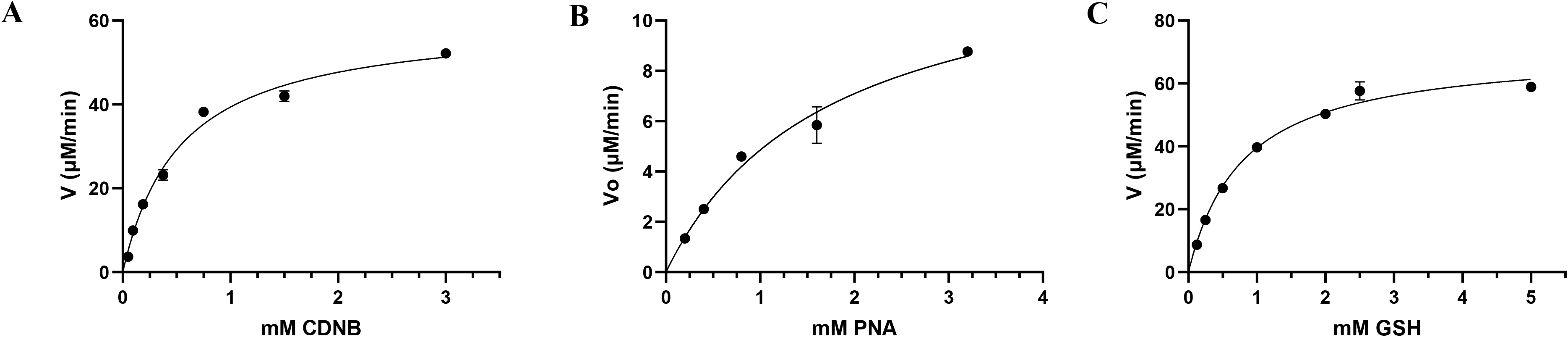
Steady-state initial velocities are plotted versus substrate concentrations. CDNB 0.05mM to 3mM. PNA concentrations were varied from 0.2mM to 3.2mM. GSH concentrations were varied from 0.125mM to 5mM. Plots were curve fit by non-linear regression and generated with GraphPad Prism.

**Table 2.** Kinetic parameters for the conjugation of GSH with CDNB and other possible substrates.

| Substrate | V<sub>max</sub><br>($\mu\text{M}/\text{min}$) | K<sub>m</sub> (mM) | K<sub>cat</sub> ($\text{min}^{-1}$) | k<sub>cat</sub>/K<sub>m</sub><br>($\text{mM}/\text{min}$) |
| --- | --- | --- | --- | --- |
| GSH | $78.2 \pm 3.46$ | $0.689 \pm 0.118$ | $44.0 \pm 1.95$ | 63.8 |
| CDNB | $60.9 \pm 3.49$ | $0.542 \pm 0.088$ | $34.1 \pm 0.63$ | 62.9 |
| PNA | $13.5 \pm 2.13$ | $1.830 \pm 0.572$ | $7.7 \pm 1.21$ | 4.2 |
| HNE | ND | ND | ND | ND |
| T2H | ND | ND | ND | ND |

**Table 3.** Currently available structures of arthropod GSTs.

### 3.4. LdGSTu1 enzyme inhibition assay

To test the interaction of LdGSTu1 with the known GST inhibitor ethacrynic acid (EA) and multiple pesticides (carbaryl, diazinon, imidacloprid, acetamiprid, chlorpyrifos, and thiamethoxam), inhibition assays were conducted by measuring change to the rate of GSH conjugation with CDNB (Fig. 6, Table 2). In the case of LdGSTu1, ethacrynic acid acted as inhibitor of GSH enzyme catalyzed conjugation to CDNB at µM concentrations, consistent with previous GST inhibition studies [60]. At a concentration of 40 µM EA, LdGSTu1 residual activity was 88.8%; at 200 µM EA, LdGSTu1 residual activity was 49.6%; and at 1mM EA, LdGSTu1 residual activity was 0.0%. Compared to EA, the inhibitory effect of the pesticides screened was relatively lower. At 40 µM, none of the pesticides showed significant inhibitory effect on the enzymatic conjugation of GSH to CDNB. However, at increasing concentrations of pesticides, the inhibitory effects became significant (Fig. 6). For the LdGSTu1, GSH catalyzed conjugation of CDNB in the presence of 1 mM acetamiprid, 1 mM carbaryl, 1 mM diazinon, 1 mM chlorpyrifos, 1 mM imidacloprid, and 1 mM thiamethoxam, the residual enzyme activity fell to 81.0%, 88.5%, 88.5%, 89.9%, 93.7%, 95.0%, respectively. In the presence of 5 mM acetamiprid, 5 mM diazinon, 5 mM chlorpyrifos, 5 mM imidacloprid, and 5 mM thiamethoxam, the residual enzyme activity was 39.1%, 75.3%, 70.5%, 66.4%, and 72.3%, respectively. Carbaryl was not included in 5 mM grouping due to insolubility and EA was not included in 5 mM grouping because at 1 mM EA residual activity already fell to 0%. These results demonstrated that the enzymatic conjugation of GSH to CDNB could be inhibited by multiple pesticides, suggesting these pesticides are potential substrates of LdGSTu1.

**Fig. 6.**
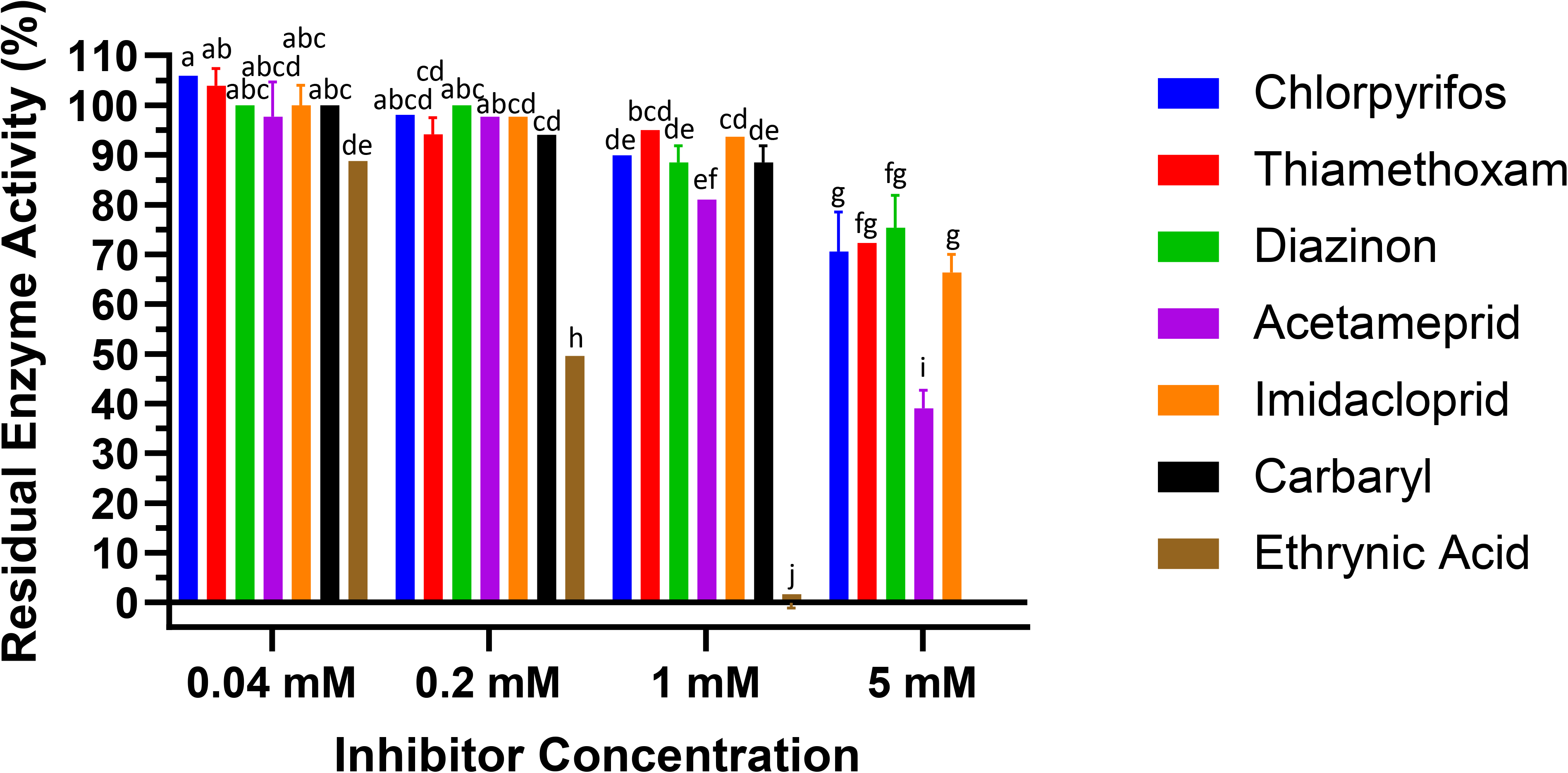
Inhibition of LdGSTu1 with ethacrynic acid (EA) and pesticides. Data points are means of independent triplicate experiments. Error bars are the calculated standard deviations of the independent triplicate experiments. Columns are color coded (see the key on right for representative colors) for the inhibitors screened and grouped into four concentrations from left to right 0.04 mM, 0.2 mM, 1 mM, 5 mM.

### 3.5 LdGSTu1 ligand docking

LdGSTu1 was docked with the ligands CDNB, EA, carbaryl, diazinon, imidacloprid, acetamiprid, and thiamethoxam to test their binding affinities. Binding poses for all ligands screened were found and ranked according to affinity dG scores. The highest ranked poses for each ligand were used for further analysis. All of the ligands docked into the presumptive hydrophobic binding site “H” of LdGSTu1 adjacent to the GSH binding site “G” and the co-crystal complexed GSH molecule with favorable calculated binding energies (Fig. S2). For CDNB, EA, carbaryl, diazinon, imidacloprid, acetamiprid, and thiamethoxam, the calculated binding energies for top poses by the affinity dG scoring function were −4.4 kcal/mol, −5.5 kcal/mol, −4.6 kcal/mol, −4.9 kcal/mol, −4.6 kcal/mol, −5.4 kcal/mol, and −5.2 kcal/mol, respectively. For the docked ligand poses with LdGSTu1, closest atom distances for ligand to GSH are listed. The CDNB ligand docked with carbon 3 positioned 3.77 Å from the glutathione sulfur atom. EA docked with carbon 2 and 3 at 4.39 Å from the glutathione sulfur atom. For carbaryl, the ligand docked with its carbonyl carbon positioned 4.01 Å from the glutathione sulfur atom. Diazinon docked with the pyrimidine ester carbon located 2.74 Å from the glutathione sulfur atom. Imidacloprid docked with C3 on the pyridine ring located 3.9 Å from the sulfur atom of glutathione. Docked acetamiprid had C7 positioned at 3.7 Å from the sulfur of GSH. Lastly, thiamethoxam docked with its sulfur atom 3.8 Å from the sulfur of GSH. Molecular docking with the crystal structure of LdGSTu1 gave favorable binding poses for all the ligands screened. Additionally, the predicted binding locations for all the highest ranked ligand poses were localized the hydrophobic binding pocket in the active site of LdGSTu1 and adjacent to the co-crystalized position of GSH (Fig. S2). The LdGSTu1 ligand docking results suggested that LdGSTu1 is capable of binding pesticides tested, further suggesting these pesticides are potential substrates of LdGSTu1.

### 3.6. mRNA expression patterns of LdGSTu1

The *LdGSTu1* expression levels between an insecticide resistant strain and the susceptible strain were examined. As shown in Fig. 7, the relative expression level of *LdGSTu1* was significantly higher in the resistant strain than the susceptible one. We then further investigated the temporal and spatial expression patterns of *LdGSTu1* in both insecticide resistant and susceptible strains. The results in Fig. 8A showed that during different developmental stages, *LdGSTu1* was expressed at the highest level during the adults (both males and females) in the insecticide resistant strain than other stages. There were significantly differences in adult expression between resistant strain and susceptible strain. The tissue expression profiles showed that *LdGST1* had the highest expression levels in the head of resistant strain, followed by midgut, Malpighian tubule, and ovary in the resistant strain. There were significantly differences in head and midgut expression between resistant strain and susceptible strain (Fig. 8B).

**Fig. 7.**
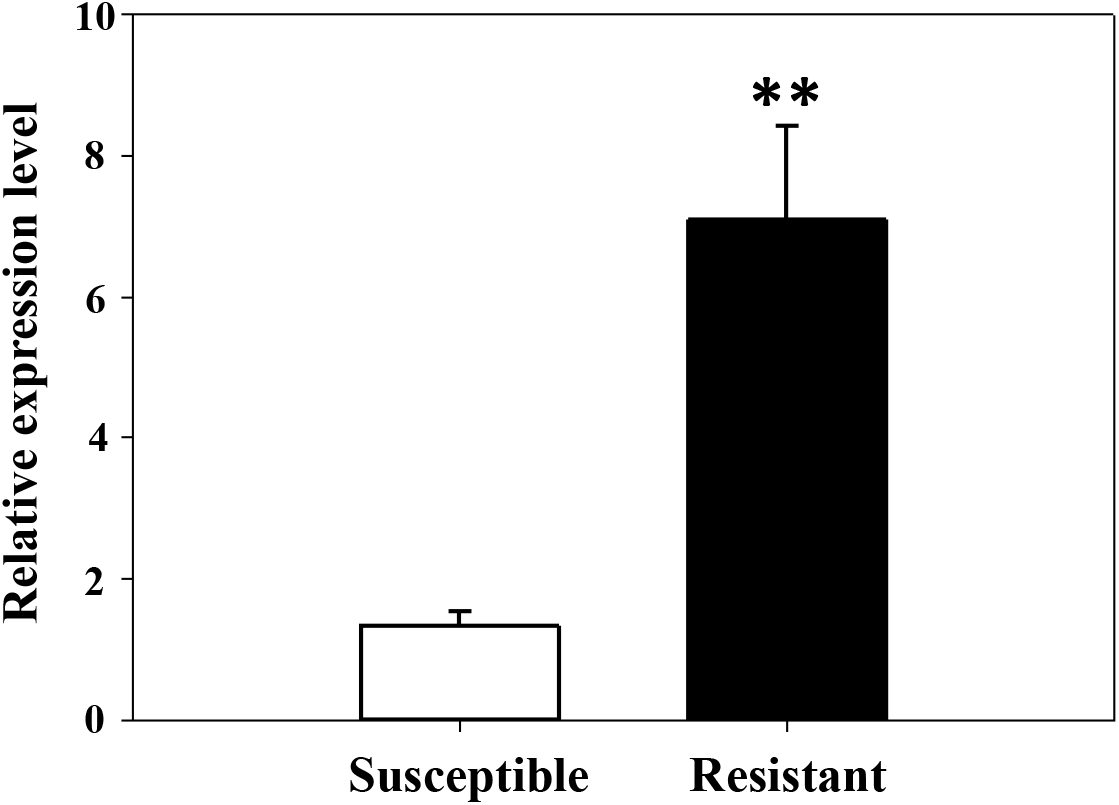
Expression patterns of *LdGSTu1* in resistant and susceptible strain. The “**” indicate significant differences in gene expression at *P* < 0.01, according to independent Student’s *t*-test.

**Fig. 8.**
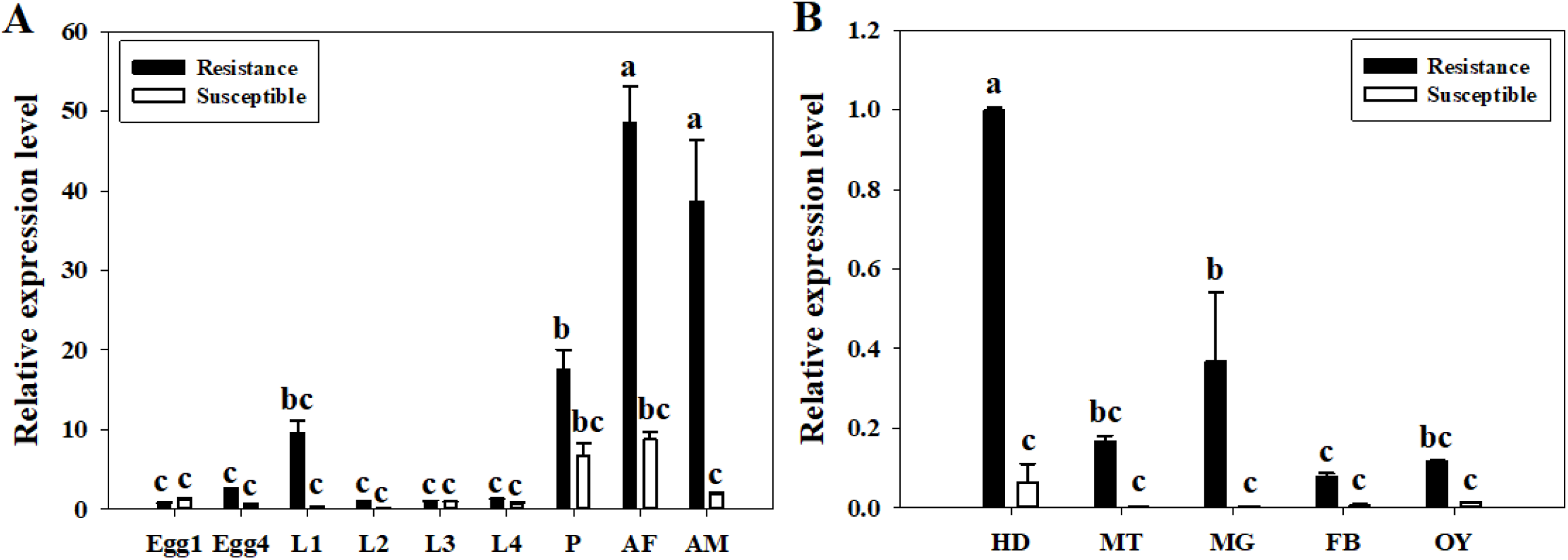
Developmental and spatial expression patterns of *LdGSTu1*. (A) Transcriptional expression levels of *LdGSTu1* during different developmental stages. Samples included: Eggs; first, second, third, and fourth instar larvae (L1-L4); pupae (P); female adults (AF); and male adults (AM). (B) Transcriptional expression levels of *LdGSTu1* in the tissues of adults. HD, heads; MT, Malpighian tubule; MG, midgut; FB, fat body; OY, ovary. Each value is the mean ± SE of three collections. Differences among multiple treatments were analyzed using one-way ANOVA, followed by a Tukey’s HSD multiple comparison test. There was no significant difference among treatments with the same alphabetic letters (e.g. a, b, and c).

## 4. Discussion

LdGSTu1 crystalized with a dimer in the asymmetric unit (Fig. S1). The dimer is a result of crystal packing and not expected to be a biologically relevant dimer assembly as the active sites are solvent exposed and not internal to the dimer interface as reported for biological dimer assemblies in previous GST crystal structure studies [17, 56, 58, 61] (Fig. S1). However, the differences between the two monomers making up the crystallographic dimer are interesting (Fig. S1). The active sites of chain A has the bound GST cofactor GSH, but chain B did not have a bound GSH molecule. Chain A complexed with GSH has an open active site similar to previously published GST structures, whereas the chain B monomer vacant of GSH has a more closed active site, suggesting flexibility in loop-helix-loop region in the active site of the N-term domain of GSTs.

Our data showed that the crystal structure of LdGSTu1 exhibited a bound GSH ligand in the “G-site” of chain A. That bound GSH revealed the hydroxyl of Ser14 to be hydrogen bonded to the thiol of GSH via a water bridge (Fig. 4A), suggesting that Ser14 is a residue responsible for catalytically activating GSH in LdGSTu1. The only other unclassified insect GST with a published crystal structure in the PDB (5ZFG) was in apo-form but also posed a crystallographic water hydrogen bonded the hydroxyl of Ser14 in BmGSTu2 [56]. Previously, characterized and classified GSTs have been shown to poses a catalytically active serine, tyrosine, or cysteine in their active sites [62]. The catalytic residue activates the glutathione thiol group through hydrogen bonding. Moreover, the unclassified insect GSTs display the sequence motif VSDGPPSL in the “G-site” which contains Ser14 (Fig. 9). In the study of BmGSTu2 (PDB: 5ZFG), the authors created a mutant P13A swapping Pro13 for an Ala [56]. It was found that the P13A mutant exhibited decreased specific activities for both CDNB and diazinon [56]. As proline residues are frequently found to be conserved in and around enzyme active sites, it can be inferred that proline residues are important for maintaining positional orientation of catalytically relevant residues. As the mutant P13A residue in BmGSTu2 directly precedes Ser14, we suggest the mutation may have caused a local structural positioning shift for Ser14, thus effecting the ability of BmGSTu2 to activate the thiol of GSH, further suggesting the catalytic importance of Ser14 in unclassified insect GSTs.

**Fig. 9.**
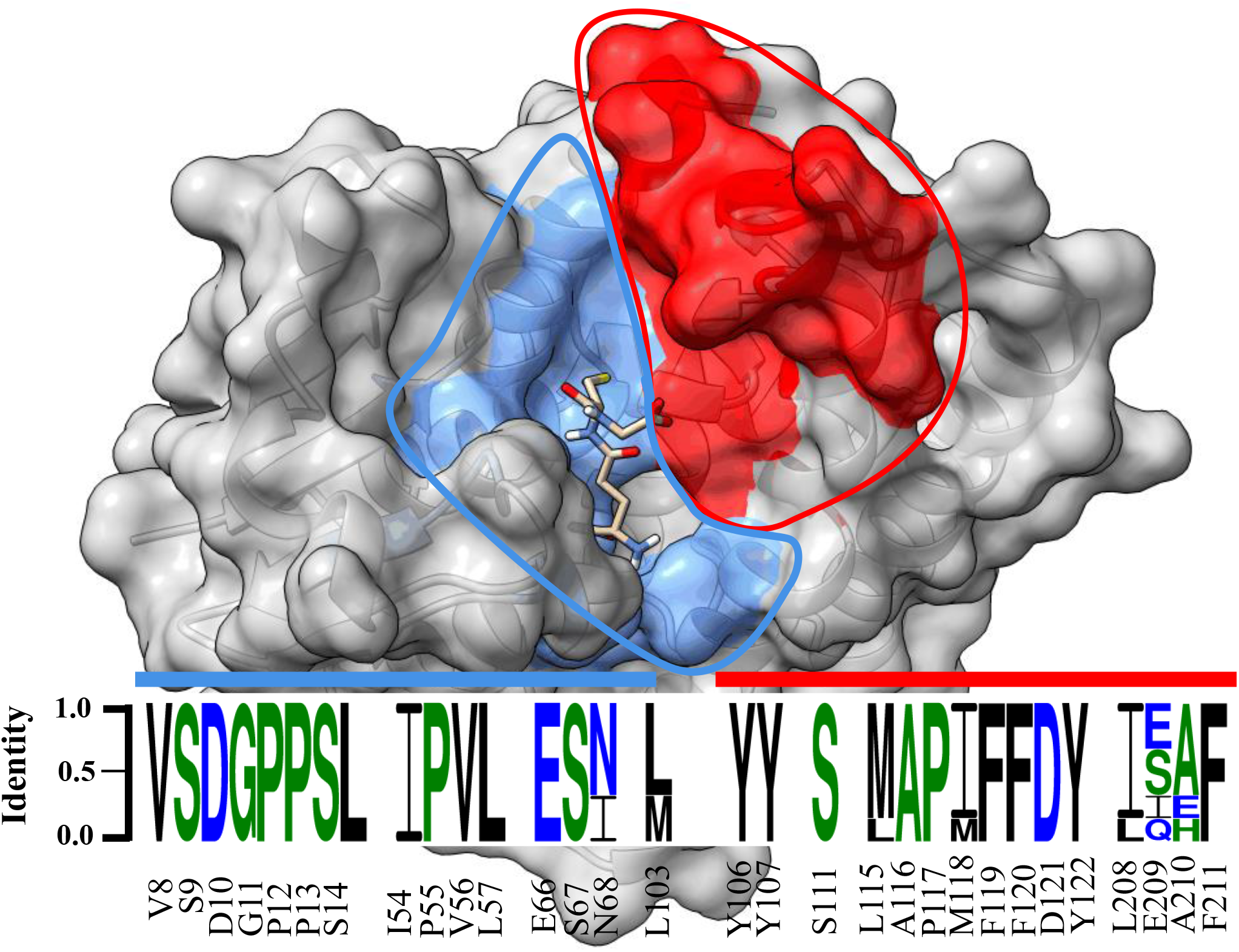
Sequence logo of amino acids in the substrate binding pocket of unclassified insect GSTs. Sequence identity is represented by the letter size. Amino acid number and type label on x-axis is representative of the LdGSTu1 sequence. The sequence logo is overlayed on a surface representation of LdGSTu1. The G-site is colored in cornflower blue and the H-site is colored in red. Residues in the sequence logo are separated by signature motifs identified in the substrate binding pocket region of LdGSTu1 upon multiple alignment with other unclassified insect GSTs. Sequences used for the multiple sequence alignment are the unclassified GST sequences from phylogenic analysis (Table S3). Sequence logo was generated with WebLogo 3 [69].

GST-mediated xenobiotic adaptation is through direct metabolism or sequestration of xenobiotics, and/or indirectly by providing protection against oxidative stress induced by xenobiotic exposure [15, 56, 57]. The GSTs being upregulated in insecticide resistant insects have been previously reported in *BmGSTu2*, which transcript expression was induced 1.7-fold in a resistant strain of *B. mori* [63]. Additionally, at the protein level, increased GST activity has been observed in insecticide resistant insects, such as an abamectin-resistant vegetable leafminer and a brown planthopper resistant to imidacloprid [64, 65]. Based on previous reports on overexpression of GSTs in insecticide resistance and increased GST activity, it was inferred that LdGSTu1 may play a role in insecticide resistance in CPB, as it is overexpressed in the insecticide resistant strain (Fig. 7). Tissue expression profile analysis showed that *LdGSTu1* expressed at the highest level in the head of resistant strain (Fig. 8B). Since the head or central nervous system is critical organ for insect survival and serves as the target for numerous neurotoxic pesticides [66, 67], the high expression of *LdGSTu1* implies its potential primary functions in xenobiotic adaptation.

Our LdGSTu1 kinetic enzyme studies showed that LdGSTu1 displayed a higher catalytic efficiency for CDNB than PNA (Table 2). LdGSTu1 enzyme inhibition assay showed that ethacrynic acid and the pesticides carbaryl, diazinon, imidacloprid, acetamiprid, chlorpyrifos, and thiamethoxam acted as inhibitors of the enzyme catalyzed conjugation of GSH to CDNB (Fig. 6, Table 2). Functional studies have previously shown insect GSTs to be associated with adaptation to plant allelochemicals and insecticides by means of direct metabolism or defense against reactive oxygen species (ROS) [15, 63]. In our study, neither HNE nor T2H were conjugated to GSH enzymatically by LdGSTu1 (Table 2). This result is consistent with bmGSTu2 and pxGSTu1, unclassified GSTs identified in silkworm [63] and diamondback moth [68], respectively.

In summary, we identified a beetle GST, LdGSTu1 belonging to the unclassified class of insect GSTs and characterized the structure and function of LdGSTu1 through x-ray crystallography, enzyme activity and binding studies. LdGSTu1 crystal structure exhibits a typical GST global fold and an active site composed of two substrate binding sites, the “G-site” and the “H-site”. The signature motif VSDGPPSL was identified, and it contains the catalytically active residue Ser14. The enzyme kinetic parameters and enzyme-substrate interaction studies demonstrated that LdGSTu1 could be inhibited by multiple pesticides tested. Further investigation is on the way to identify putative catalytic active residues through site-directed mutagenesis along with continuing enzyme activity and binding studies to identify if LdGSTu1 only binds pesticides or whether it can also metabolize them.

## Acknowledgments

This work was supported by a faculty start-up fund from Pennsylvania State University, and the USDA National Institute of Food and Federal Appropriations under Hatch Project #PEN04609 and Accession #1010058. Yanjun Liu was supported by a fellowship from the China Scholarship Council and the earmarked fund for China Agriculture Research System [grant number CARS-28]. Timothy Moural was supported by USDA NIFA postdoctoral fellowship.

## Author contributions

F.Z. and T.M. conceived and designed the experiments. Y.L., S.K., BK, J.H., Z.S., T.M. conducted bioinformatics, molecular biology, and biochemistry studies. T.M. and Y.L. conducted protein crystallography studies. F.Z., T.M., A.A. contributed new reagents/analytic tools. Y.L., T.M., F.Z. wrote the original draft; all authors contributed to revise and prepare the paper for submission.

## Supplementary Information

**Fig. S1.**
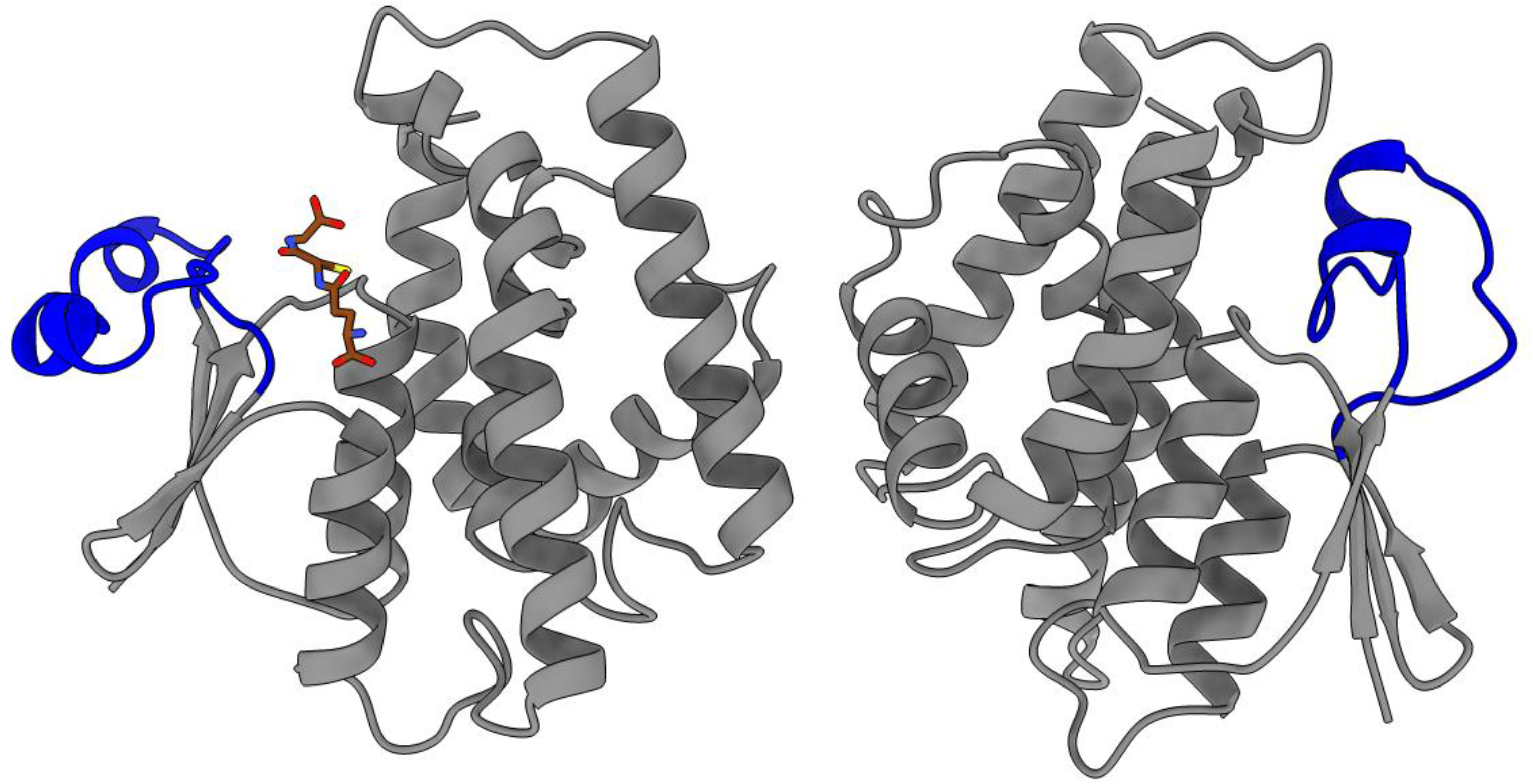
LdGSTu1 crystallographic dimer. Chain A exhibited glutathione (GSH) bound to the active site (left monomer). Chain B had no bound GSH molecule (right monomer). Monomers are depicted in gray with the loop helices loop region of N-term domain in blue. GSH is colored by heteroatom with carbon colored brown.

**Fig. S2.**
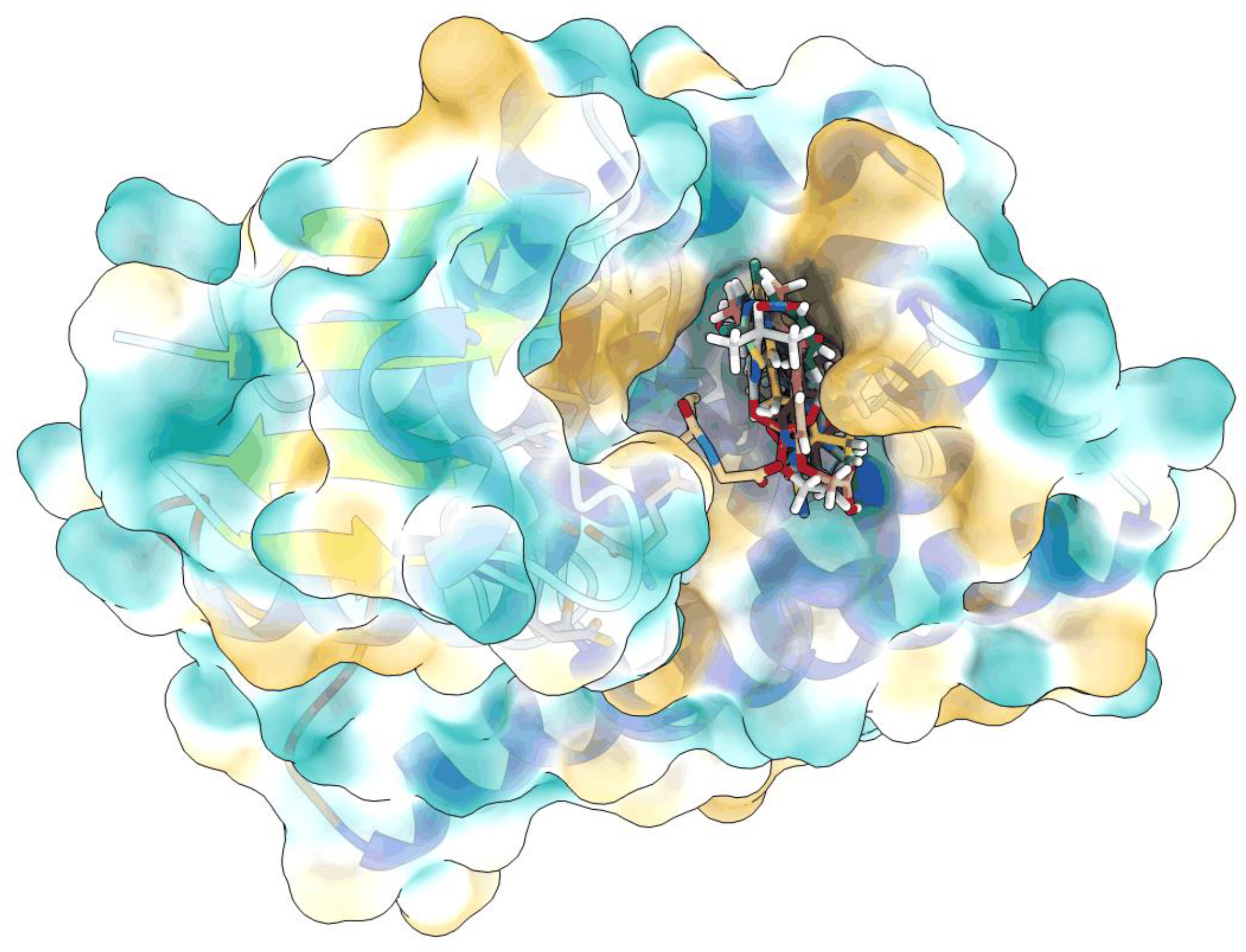
LdGSTu1 was docked with the ligands CDNB, EA, carbaryl, diazinon, imidacloprid, acetamiprid, and thiamethoxam. The highest ranked docked poses are shown bound in the LdGSTu1 (surface overlayed on ribbon diagram and colored by hydrophobicity) active site. Ligands are shown as stick models colored by heteroatom with carbons colored according to ligand. For docked ligands on the right the carbons are colored in light blue, pink, forest green, light gray, purple, yellow, and light green for CDNB, EA, carbaryl, diazinon, imidacloprid, acetamiprid, and thiamethoxam, respectively. The co-crystalized GSH is shown on the left of the active site, represented in stick format and colored by heteroatom with carbon atoms colored tan.

**Table S1.** Currently available structures of insect GSTs.

| Order | Species | PDB ID | GST name | Ligand | Resolution (Å) | Reference | No. |
| --- | --- | --- | --- | --- | --- | --- | --- |
| Diptera | *Anopheles dirus* | 1JLV | Delta 3 | GSH | 1.75 Å | Oakely et al., 2001 | 1 |
|  | *Anopheles dirus* | 1JLW | Delta 4 | Apo | 2.45 Å | Oakely et al., 2001 | 2 |
|  | *Anopheles dirus* | 3F63 | Delta 4 | GTX | 1.80 Å | Wongsantichon et al, 2010 | 3 |
|  | *Anopheles dirus* | 3F6D | Delta 4 F123A | GTX | 1.70 Å | Wongsantichon et al, 2010 | 4 |
|  | *Anopheles dirus* | 3G7I | Delta 4 | Apo and GSW | 2.05 Å | Wongsantichon et al, 2010 | 5 |
|  | *Anopheles dirus* | 3G7J | Delta 4 Y119E | GTX | 2.20 Å | Wongsantichon et al, 2010 | 6 |
|  | *Anopheles dirus* | 1R5A | Delta 5 | CU | 2.50 Å | Udomsinprasert et al., 2005 | 7 |
|  | *Anopheles dirus* | 1V2A | Delta 6 | GTS | 2.15 Å | Udomsinprasert et al., 2005 | 8 |
|  | *Anopheles gambiae* | 1PN9 | Delta 1-6 | GTX | 2.00 Å | Chen et al., 2003 | 9 |
|  | *Anopheles gambiae* | 2IL3 | Epsilon 2 | Apo | 2.20 Å | Wang et al., 2008 | 10 |
|  | *Anopheles gambiae* | 2IMI | Epsilon 2 | GSH | 1.40 Å | Wang et al., 2008 | 11 |
|  | *Anopheles gambiae* | 2IMK | Epsilon 2 | GTX | 1.90 Å | Wang et al., 2008 | 12 |
|  | *Anopheles gambiae* | 4GSN | Epsilon 2 | 1PE; GOL; GSH | 2.30 Å | Mitchell et al., 2014 | 13 |
|  | *Drosophila melanogaster* | 3EIN | Delta 1 | GTT | 1.13 Å | Low et al., 2010 | 14 |
|  | *Drosophila melanogaster* | 5F0G | Delta 2 | K<sup>+</sup>; Na<sup>+</sup> | 1.6 Å | Gonzalez, et al., 2018 | 15 |
|  | *Drosophila melanogaster* | 3F6F | Delta 10 | Apo | 1.60 Å | Wongsantichon et al, 2010 | 16 |
|  | *Drosophila melanogaster* | 3GH6 | Delta 10 | GTT | 1.65 Å | Wongsantichon et al, 2010 | 17 |
|  | *Drosophila melanogaster* | 1MOU | Sigma 1 | GSW | 1.75 Å | Singh et al. 2001; Agianian et al., 2003 | 18 |
|  | *Drosophila melanogaster* | 4PNF | Epsilon 6 | GSH | 2.11 Å | Scian et al., 2015 | 19 |
|  | *Drosophila melanogaster* | 4PNG | Epsilon 7 | GSF | 1.53 Å | Scian et al., 2015 | 20 |
|  | *Drosophila melanogaster* | 7DB0 | Epsilon 14 | DC1; DMS | 1.66 Å | Koiwai et al., 2021 | 21 |
|  | *Scaptomyza nigrita* | 4I97 | Delta 1 | GSH | 2.15 Å | Gloss et al., 2014 | 22 |
|  | *Musca domestica* | 5ZWP | Delta 1 | - | 1.40 Å | Sue and Yajima, 2018 | 23 |
|  | *Musca domestica* | 3VWX | Epsilon | GSH | 1.80 Å | Nakamura et al., 2013 | 24 |
| Lepidoptera | *Bombyx mori* | 3VK9 | Delta | - | 2.00 Å | Yamamoto et al., 2012 | 25 |
|  | *Bombyx mori* | 3WD6 | Omega | - | 2.50 Å | Yamamoto et al., 2013 | 26 |
|  | *Bombyx mori* | 3AY8 | Unclassified | - | 2.10 Å | Yamamoto et al., 2011 | 27 |
|  | *Bombyx mori* | 5ZFG | Unclassified 2 | - | 1.70 Å | Yamamoto et al., 2018 | 28 |
|  | *Nilaparvata lugens* | 3WYW | Delta | - | 1.70 Å | Yamamoto et al., 2015 | 29 |
| Coleoptera | *Leptinotarsa decemlineata* | 7RKA | Unclassified 1 | GSH | 1.80 Å | This study | 30 |

**Table S2.**
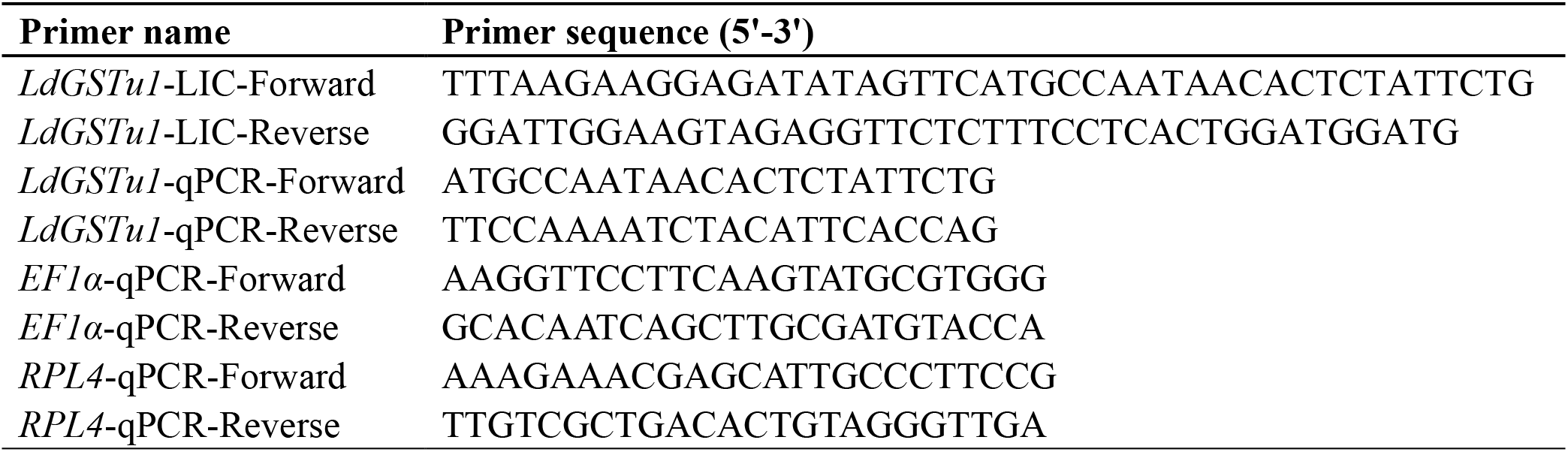
Primers used in the current study.

**Table S3.** Protein information of GSTs used for phylogenetic analysis.

| **Insect species** | **GST name** | **Accession number** | **Length (aa)** | **Reference** |
| --- | --- | --- | --- | --- |
| *Anopheles cracens* | AcGSTo1 | ACY_95464.1 | 248 | Wongtrakul et al. 2010 JME |
| *Anopheles dirus* | AdGSTe1 | ABD_43203.1 | 221 |  |
| *Anopheles gambiae* | AgGSTe1 | AAL_59658.1 | 224 | Ortelli et al. 2003. Biochem. J |
|  | AgGSTz1 | AAM61889.1 | 222 | Ding et al. 2003. BMC Genomics |
| *Anoplophora glabripennis* | AgGST1-1 | XP_018564199.1 | 231 |  |
| *Apis mellifera* | AmGSTd1 | NP_001171499.1 | 217 | Elsik et al. 2014. BMC Genomics |
|  | AmGSTt3 | XP_624692.2 | 230 |  |
|  | AmGSTs4 | NP_001136128.1 | 206 | Elsik et al. 2014. BMC Genomics |
| *Bombyx mori* | BmGSTe4 | NP_001108460.1 | 217 | Yamamoto et al. 2013 Insect Mol Biol |
|  | BmGSTu2 | NP_001108462.1 | 233 | Yu et al. 2008 Insect Biochem Mol Biol |
|  | BmGSTt1 | NP_001108463.1 | 229 | Yu et al. 2008 Insect Biochem Mol Biol |
|  | BmGSTz1 | NP_001037418.1 | 215 | Yu et al. 2008 Insect Biochem Mol Biol |
|  | BmGSTs1 | NP_001037077.1 | 206 | Yu et al. 2008 Insect Biochem Mol Biol |
| *Drosophila melanogaster* | DmGSTe1 | NP_611323.1 | 224 |  |
|  | DmGSTd6 | NP_524915.1 | 215 | Sun et al. 2006 IMB |
|  | DmGSTo1 | NP_648237.1 | 254 |  |
| *Drosophila mauritiana* | DmGST1-1-X2 | XP_033167954 | 265 |  |
| *Drosophila mojavensis* | DmGST1-X2 | XP_015022387.1 | 231 |  |
| *Leptinotarsa decemlineata* | LdGSTe1 | APX61028.1 | 217 | Han et al. 2016 Pesticide Biochem Physiol |
|  | LdGSTd3 | APX61027.1 | 216 | Han et al. 2016 Pesticide Biochem Physiol |
|  | LdGSTu1 | XP_023027125.1 | 230 | This study |
|  | LdGSTt2 | APX61052.1 | 224 | Han et al. 2016 Pesticide Biochem Physiol |
|  | LdGSTs1 | APX61045.1 | 204 | Han et al. 2016 Pesticide Biochem Physiol |
| *Locusta migratoria* | LmGSTs1 | AEB91973.1 | 204 | Qin et al. 2011 Pest Manag Sci |
|  | LmGSTs2 | AEB91974.1 | 204 | Qin et al. 2011 Pest Manag Sci |
| *Nasonia vitripennis* | NvGSTd1 | NP_001165913.1 | 215 | Oakeshott et al. 2010 Insect Mol Biol |
|  | NvGSTt3 | NP_001165927.1 | 221 | Oakeshott et al. 2010 Insect Mol Biol |
|  | NvGSTo1 | NP_001165912.1 | 241 | Oakeshott et al. 2010 Insect Mol Biol |
|  | NvGSTz1 | NP_001165931.1 | 217 | Oakeshott et al. 2010 Insect Mol Biol |
| *Spodoptera litura* | SIGSTz2 | AIH07599.1 | 212 |  |
| *Sitophilus oryzae* | SoGST1-X2 | XP_030760484.1 | 239 |  |

## References

[1] A.A. Enayati, H. Ranson, J. Hemingway, Insect glutathione transferases and insecticide resistance, Insect molecular biology 14(1) (2005) 3–8.

[2] R. Friedman, Genomic organization of the glutathione S-transferase family in insects, Molecular phylogenetics and evolution 61(3) (2011) 924–32.

[3] C. Frova, Glutathione transferases in the genomics era: new insights and perspectives, Biomol Eng 23(4) (2006) 149–69.

[4] D. Sheehan, G. Meade, V.M. Foley, C.A. Dowd, Structure, function and evolution of glutathione transferases: implications for classification of non-mammalian members of an ancient enzyme superfamily, The Biochemical journal 360(Pt 1) (2001) 1–16.

[5] R.N. Armstrong, Glutathione S-transferases: reaction mechanism, structure, and function, Chemical research in toxicology 4(2) (1991) 131–40.

[6] J.D. Hayes, J.U. Flanagan, I.R. Jowsey, Glutathione transferases, Annual review of pharmacology and toxicology 45 (2005) 51–88.

[7] N. Lumjuan, B.J. Stevenson, L.A. Prapanthadara, P. Somboon, P.M. Brophy, B.J. Loftus, D.W. Severson, H. Ranson, The *Aedes aegypti* glutathione transferase family, Insect biochemistry and molecular biology 37(10) (2007) 1026–35.

[8] N. Allocati, L. Federici, M. Masulli, C. Di Ilio, Glutathione transferases in bacteria, The FEBS journal 276(1) (2009) 58–75.

[9] R. Friedman, Genomic organization of the glutathione S-transferase family in insects, Molecular Phylogenetics and Evolution 61(3) (2011) 924–932.

[10] H. Shi, L. Pei, S. Gu, S. Zhu, Y. Wang, Y. Zhang, B. Li, Glutathione S-transferase (GST) genes in the red flour beetle, *Tribolium castaneum*, and comparative analysis with five additional insects, Genomics 100(5) (2012) 327–335.

[11] H. Ranson, C. Claudianos, F. Ortelli, C. Abgrall, J. Hemingway, M.V. Sharakhova, M.F. Unger, F.H. Collins, R. Feyereisen, Evolution of supergene families associated with insecticide resistance, Science (New York, N.Y.) 298(5591) (2002) 179–81.

[12] J.M. Riveron, C. Yunta, S.S. Ibrahim, R. Djouaka, H. Irving, B.D. Menze, H.M. Ismail, J. Hemingway, H. Ranson, A. Albert, C.S. Wondji, A single mutation in the GSTe2 gene allows tracking of metabolically based insecticide resistance in a major malaria vector, Genome Biol 15(2) (2014) R27.

[13] A.J. Ketterman, C. Saisawang, J. Wongsantichon, Insect glutathione transferases, Drug metabolism reviews 43(2) (2011) 253–65.

[14] A.J. Oakley, Glutathione transferases: new functions, Current opinion in structural biology 15(6) (2005) 716–23.

[15] N. Pavlidi, J. Vontas, T. Van Leeuwen, The role of glutathione S-transferases (GSTs) in insecticide resistance in crop pests and disease vectors, Curr Opin Insect Sci 27 (2018) 97–102.

[16] L.W. Meng, G.R. Yuan, X.P. Lu, T.X. Jing, L.S. Zheng, H.X. Yong, J.J. Wang, Two delta class glutathione S-transferases involved in the detoxification of malathion in *Bactrocera dorsalis* (Hendel), Pest management science 75(6) (2019) 1527–1538.

[17] D. Gonzalez, S. Fraichard, P. Grassein, P. Delarue, P. Senet, A. Nicolaï, E. Chavanne, E. Mucher, Y. Artur, J.F. Ferveur, J.M. Heydel, L. Briand, F. Neiers, Characterization of a *Drosophila* glutathione transferase involved in isothiocyanate detoxification, Insect biochemistry and molecular biology 95 (2018) 33–43.

[18] H. Yan, H. Jia, H. Gao, X. Guo, B. Xu, Identification, genomic organization, and oxidative stress response of a sigma class glutathione S-transferase gene (AccGSTS1) in the honey bee, *Apis cerana cerana*, Cell stress & chaperones 18(4) (2013) 415–26.

[19] H. Yan, H. Jia, X. Wang, H. Gao, X. Guo, B. Xu, Identification and characterization of an *Apis cerana cerana* Delta class glutathione S-transferase gene (AccGSTD) in response to thermal stress, Die Naturwissenschaften 100(2) (2013) 153–63.

[20] X. Zou, Z. Xu, H. Zou, J. Liu, S. Chen, Q. Feng, S. Zheng, Glutathione S-transferase SlGSTE1 in *Spodoptera litura* may be associated with feeding adaptation of host plants, Insect biochemistry and molecular biology 70 (2016) 32–43.

[21] Y. Ding, F. Ortelli, L.C. Rossiter, J. Hemingway, H. Ranson, The *Anopheles gambiae* glutathione transferase supergene family: annotation, phylogeny and expression profiles, BMC genomics 4(1) (2003) 35.

[22] C. Claudianos, H. Ranson, R.M. Johnson, S. Biswas, M.A. Schuler, M.R. Berenbaum, R. Feyereisen, J.G. Oakeshott, A deficit of detoxification enzymes: pesticide sensitivity and environmental response in the honeybee, Insect molecular biology 15(5) (2006) 615–36.

[23] D.D. McKenna, S. Shin, D. Ahrens, M. Balke, C. Beza-Beza, D.J. Clarke, A. Donath, H.E. Escalona, F. Friedrich, H. Letsch, S. Liu, D. Maddison, C. Mayer, B. Misof, P.J. Murin, O. Niehuis, R.S. Peters, L. Podsiadlowski, H. Pohl, E.D. Scully, E.V. Yan, X. Zhou, A. Ślipiński, R.G. Beutel, The evolution and genomic basis of beetle diversity, Proceedings of the National Academy of Sciences of the United States of America 116(49) (2019) 24729–24737.

[24] A. Alyokhin, M. Baker, D. Mota-Sanchez, G. Dively, E. Grafius, Colorado Potato Beetle Resistance to Insecticides, American Journal of Potato Research 85(6) (2008) 395–413.

[25] D. Weber, Colorado beetle: Pest on the move., Pesticide Outlook 14 (2003) 256–259.

[26] A. Alyokhin, Y.H. Chen, Adaptation to toxic hosts as a factor in the evolution of insecticide resistance, Curr Opin Insect Sci 21 (2017) 33–38.

[27] S.D. Schoville, Y.H. Chen, M.N. Andersson, J.B. Benoit, A. Bhandari, J.H. Bowsher, K. Brevik, K. Cappelle, M.M. Chen, A.K. Childers, C. Childers, O. Christiaens, J. Clements, E.M. Didion, E.N. Elpidina, P. Engsontia, M. Friedrich, I. Garcia-Robles, R.A. Gibbs, C. Goswami, A. Grapputo, K. Gruden, M. Grynberg, B. Henrissat, E.C. Jennings, J.W. Jones, M. Kalsi, S.A. Khan, A. Kumar, F. Li, V. Lombard, X. Ma, A. Martynov, N.J. Miller, R.F. Mitchell, M. Munoz-Torres, A. Muszewska, B. Oppert, S.R. Palli, K.A. Panfilio, Y. Pauchet, L.C. Perkin, M. Petek, M.F. Poelchau, E. Record, J.P. Rinehart, H.M. Robertson, A.J. Rosendale, V.M. Ruiz-Arroyo, G. Smagghe, Z. Szendrei, G.W.C. Thomas, A.S. Torson, I.M. Vargas Jentzsch, M.T. Weirauch, A.D. Yates, G.D. Yocum, J.S. Yoon, S. Richards, A model species for agricultural pest genomics: the genome of the Colorado potato beetle, *Leptinotarsa decemlineata* (Coleoptera: Chrysomelidae), Scientific reports 8(1) (2018) 1931.

[28] J.A. Argentine, K.Y. Zhu, S.H. Lee, J.M. Clark, Biochemical-mechanisms of azinphosmethyl resistance in isogenic strains of Colorado potato beetle, Pesticide Biochemistry and Physiology 48(1) (1994) 63–78.

[29] K.Y. Zhu, J.M. Clark, Validation of a point mutation of acetylcholinesterase in Colorado potato beetle by polymerase chain reaction coupled to enzyme inhibition assay, Pesticide Biochemistry and Physiology 57(1) (1997) 28–35.

[30] N. Liu, F. Zhu, Q. Xu, J.W. Pridgeon, X. Gao, Behavioral change, physiological modification, and metabolic detoxification: mechanisms of insecticide resistance, Acta Entomologica Sinica 49 (2006) 671–679.

[31] F. Zhu, T.W. Moural, D.R. Nelson, S.R. Palli, A specialist herbivore pest adaptation to xenobiotics through up-regulation of multiple Cytochrome P450s, Scientific reports 6 (2016) 20421.

[32] X. Li, M.A. Schuler, M.R. Berenbaum, Molecular mechanisms of metabolic resistance to synthetic and natural xenobiotics, Annual review of entomology 52 (2007) 231–53.

[33] W. Dermauw, N. Wybouw, S. Rombauts, B. Menten, J. Vontas, M. Grbic, R.M. Clark, R. Feyereisen, T. Van Leeuwen, A link between host plant adaptation and pesticide resistance in the polyphagous spider mite *Tetranychus urticae*, Proceedings of the National Academy of Sciences of the United States of America 110(2) (2013) E113–E122.

[34] A.W. Adesanya, M.J. Beauchamp, M.D. Lavine, L.C. Lavine, F. Zhu, D.B. Walsh, Physiological resistance alters behavioral response of *Tetranychus urticae* to acaricides, Scientific reports 9(1) (2019) 19308.

[35] J. Clements, S. Schoville, N. Peterson, A.S. Huseth, Q. Lan, R.L. Groves, RNA interference of three up-regulated transcripts associated with insecticide resistance in an imidacloprid resistant population of *Leptinotarsa decemlineata*, Pestic Biochem Physiol 135 (2017) 35–40.

[36] J.B. Han, G.Q. Li, P.J. Wan, T.T. Zhu, Q.W. Meng, Identification of glutathione S-transferase genes in *Leptinotarsa decemlineata* and their expression patterns under stress of three insecticides, Pestic Biochem Physiol 133 (2016) 26–34.

[37] F. Zhu, Y. Cui, D.B. Walsh, L.C. Lavine, Application of RNAi towards insecticide resistance management, in: R. Chandrasekar, B.K. Tyagi, Z. Gui, G.R. Reeck (Eds.), Short Views on Insect Biochemistry and Molecular Biology, Academic Publisher, Manhattan, USA, 2014, pp. 595–619.

[38] C. Aslanidis, P.J. de Jong, Ligation-independent cloning of PCR products (LIC-PCR), Nucleic acids research 18(20) (1990) 6069–74.

[39] R.C. Edgar, MUSCLE: multiple sequence alignment with high accuracy and high throughput, Nucleic acids research 32(5) (2004) 1792–1797.

[40] S. Kumar, G. Stecher, M. Li, C. Knyaz, K. Tamura, MEGA X: Molecular Evolutionary Genetics Analysis across Computing Platforms, Molecular biology and evolution 35(6) (2018) 1547–1549.

[41] Database resources of the National Center for Biotechnology Information, Nucleic acids research 46(D1) (2018) D8–d13.

[42] W. Kabsch, XDS, Acta Crystallogr D Biol Crystallogr 66(Pt 2) (2010) 125–132.

[43] D. Liebschner, P.V. Afonine, M.L. Baker, G. Bunkóczi, V.B. Chen, T.I. Croll, B. Hintze, L.W. Hung, S. Jain, A.J. McCoy, N.W. Moriarty, R.D. Oeffner, B.K. Poon, M.G. Prisant, R.J. Read, J.S. Richardson, D.C. Richardson, M.D. Sammito, O.V. Sobolev, D.H. Stockwell, T.C. Terwilliger, A.G. Urzhumtsev, L.L. Videau, C.J. Williams, P.D. Adams, Macromolecular structure determination using X-rays, neutrons and electrons: recent developments in Phenix, Acta crystallographica. Section D, Structural biology 75(Pt 10) (2019) 861–877.

[44] P. Emsley, K. Cowtan, Coot: model-building tools for molecular graphics, Acta Crystallogr D Biol Crystallogr 60(Pt 12 Pt 1) (2004) 2126–32.

[45] P. Emsley, B. Lohkamp, W.G. Scott, K. Cowtan, Features and development of Coot, Acta Crystallogr D Biol Crystallogr 66(Pt 4) (2010) 486–501.

[46] S.F. Altschul, W. Gish, W. Miller, E.W. Myers, D.J. Lipman, Basic local alignment search tool, Journal of molecular biology 215(3) (1990) 403–10.

[47] H.M. Berman, J. Westbrook, Z. Feng, G. Gilliland, T.N. Bhat, H. Weissig, I.N. Shindyalov, P.E. Bourne, The Protein Data Bank, Nucleic acids research 28(1) (2000) 235–242.

[48] E.F. Pettersen, T.D. Goddard, C.C. Huang, G.S. Couch, D.M. Greenblatt, E.C. Meng, T.E. Ferrin, UCSF Chimera--a visualization system for exploratory research and analysis, Journal of computational chemistry 25(13) (2004) 1605–12.

[49] E.F. Pettersen, T.D. Goddard, C.C. Huang, E.C. Meng, G.S. Couch, T.I. Croll, J.H. Morris, T.E. Ferrin, UCSF ChimeraX: Structure visualization for researchers, educators, and developers, Protein science : a publication of the Protein Society 30(1) (2021) 70–82.

[50] T.D. Goddard, C.C. Huang, E.C. Meng, E.F. Pettersen, G.S. Couch, J.H. Morris, T.E. Ferrin, UCSF ChimeraX: Meeting modern challenges in visualization and analysis, Protein science : a publication of the Protein Society 27(1) (2018) 14–25.

[51] W.H. Habig, M.J. Pabst, W.B. Jakoby, Glutathione S-transferases. The first enzymatic step in mercapturic acid formation, The Journal of biological chemistry 249(22) (1974) 7130–9.

[52] W.H. Habig, W.B. Jakoby, [51] Assays for differentiation of glutathione S-Transferases, Methods in Enzymology, Academic Press 1981, pp. 398–405.

[53] Molecular Operating Environment (MOE), 2020.09, Chemical Computing Group ULC, 1010 Sherbrooke St. West, Suite #910, Montreal, QC, Canada, H3A 2R7, 2020.

[54] S. Kim, J. Chen, T. Cheng, A. Gindulyte, J. He, S. He, Q. Li, B.A. Shoemaker, P.A. Thiessen, B. Yu, L. Zaslavsky, J. Zhang, E.E. Bolton, PubChem in 2021: new data content and improved web interfaces, Nucleic acids research 49(D1) (2021) D1388–D1395.

[55] E.F. Pettersen, T.D. Goddard, C.C. Huang, E.C. Meng, G.S. Couch, T.I. Croll, J.H. Morris, T.E. Ferrin, UCSF ChimeraX: Structure visualization for researchers, educators, and developers, Protein Science 30(1) (2021) 70–82.

[56] K. Yamamoto, A. Higashiura, A. Hirowatari, N. Yamada, T. Tsubota, H. Sezutsu, A. Nakagawa, Characterisation of a diazinon-metabolising glutathione S-transferase in the silkworm *Bombyx mori* by X-ray crystallography and genome editing analysis, Scientific reports 8(1) (2018) 16835.

[57] L. Chen, P.R. Hall, X.E. Zhou, H. Ranson, J. Hemingway, E.J. Meehan, Structure of an insect delta-class glutathione S-transferase from a DDT-resistant strain of the malaria vector *Anopheles gambiae*, Acta crystallographica. Section D, Biological crystallography 59(Pt 12) (2003) 2211–7.

[58] A.J. Oakley, T. Harnnoi, R. Udomsinprasert, K. Jirajaroenrat, A.J. Ketterman, M.C. Wilce, The crystal structures of glutathione S-transferases isozymes 1-3 and 1-4 from *Anopheles dirus species* B, Protein science : a publication of the Protein Society 10(11) (2001) 2176–85.

[59] K. Yamamoto, A. Higashiura, M.T. Hossain, N. Yamada, T. Shiotsuki, A. Nakagawa, Structural characterization of the catalytic site of a *Nilaparvata lugens* delta-class glutathione transferase, Archives of biochemistry and biophysics 566 (2015) 36–42.

[60] G. Qin, M. Jia, T. Liu, X. Zhang, Y. Guo, K.Y. Zhu, E. Ma, J. Zhang, Characterization and functional analysis of four glutathione S-transferases from the migratory locust, *Locusta migratoria*, PloS one 8(3) (2013) e58410.

[61] Y. Mathieu, P. Prosper, M. Buée, S. Dumarçay, F. Favier, E. Gelhaye, P. Gérardin, L. Harvengt, J.P. Jacquot, T. Lamant, E. Meux, S. Mathiot, C. Didierjean, M. Morel, Characterization of a *Phanerochaete chrysosporium* glutathione transferase reveals a novel structural and functional class with ligandin properties, The Journal of biological chemistry 287(46) (2012) 39001–11.

[62] N. Allocati, M. Masulli, C. Di Ilio, L. Federici, Glutathione transferases: substrates, inihibitors and pro-drugs in cancer and neurodegenerative diseases, Oncogenesis 7(1) (2018) 8.

[63] K. Yamamoto, N. Yamada, Identification of a diazinon-metabolizing glutathione S-transferase in the silkworm, *Bombyx mori*, Scientific reports 6(1) (2016) 30073.

[64] V.M. Malathi, S.K. Jalali, D.K. Gowda, M. Mohan, T. Venkatesan, Establishing the role of detoxifying enzymes in field-evolved resistance to various insecticides in the brown planthopper (*Nilaparvata lugens*) in South India, Insect science 24(1) (2017) 35–46.

[65] Q.B. Wei, Z.R. Lei, R. Nauen, D.C. Cai, Y.L. Gao, Abamectin resistance in strains of vegetable leafminer, *Liriomyza sativae* (Diptera: Agromyzidae) is linked to elevated glutathione S-transferase activity, Insect science 22(2) (2015) 243–50.

[66] F. Zhu, R. Parthasarathy, H. Bai, K. Woithe, M. Kaussmann, R. Nauen, D.A. Harrison, S.R. Palli, A brain-specific cytochrome P450 responsible for the majority of deltamethrin resistance in the QTC279 strain of *Tribolium castaneum*, Proceedings of the National Academy of Sciences of the United States of America 107(19) (2010) 8557–62.

[67] C. Manjon, B.J. Troczka, M. Zaworra, K. Beadle, E. Randall, G. Hertlein, K.S. Singh, C.T. Zimmer, R.A. Homem, B. Lueke, R. Reid, L. Kor, M. Kohler, J. Benting, M.S. Williamson, T.G.E. Davies, L.M. Field, C. Bass, R. Nauen, Unravelling the molecular determinants of bee sensitivity to neonicotinoid insecticides, Current biology : CB 28(7) (2018) 1137–1143.e5.

[68] K. Yamamoto, A. Hirowatari, T. Shiotsuki, N. Yamada, Biochemical characterization of an unclassified glutathione S-transferase of *Plutella xylostella*, Journal of pesticide science 41(4) (2016) 145–151.

[69] G.E. Crooks, G. Hon, J.M. Chandonia, S.E. Brenner, WebLogo: a sequence logo generator, Genome research 14(6) (2004) 1188–90.

